# Elusive tropical forest canopy diversity revealed through environmental DNA contained in rainwater

**DOI:** 10.1101/2025.02.26.640397

**Authors:** Lucie Zinger, Anne-Sophie Benoiston, Yves Cuenot, Céline Leroy, Eliane Louisanna, Lucie Moreau, Frédéric Petitclerc, Finn Piatscheck, Jérôme Orivel, Cécile Richard-Hansen, Lou Hansen-Chaffard, Uxue Suescun, Valérie Troispoux, Frédéric Boyer, Jérôme Chave, Thibaud Decaëns, Antoine Fouquet, Johan Pansu, Jérémy Raynaud, Rodolphe Rougerie, Lucas Sire, Heidy Schimann, Pierre Taberlet, Amaia Iribar

## Abstract

Tropical rainforest canopies shelter an under-explored reservoir of biodiversity. In recent years, the amplification and sequencing of taxonomically informative DNA fragments from environmental samples (i.e. eDNA) has revolutionized biomonitoring. Here, we explore the potential of DNA contained in canopy throughfall water to sample the biological diversity of rainforest canopies. By sampling rainwash eDNA in two 1ha-plots, one mature Amazonian forest and a nearby tree plantation, we were able to detect 170 plant taxa, 72 vertebrate taxa including mammals, birds, and amphibians, and 313 insect taxa including e.g. mosquitoes, ants, beetles. The taxonomic composition retrieved in these two plots reflected their different disturbance status, and the plantation exhibited a much lower diversity. We also found that rainwash eDNA persists over ten days in passive eDNA collectors while providing a local picture of the diversity. This suggests that this method can be scaled up for a cost-effective environmental management of rainforest’, and more generally of all forests canopies.

**Teaser:** Rainwash DNA reveals the identity of plants, animals, and insects diversity, offering a practical tool for monitoring canopy biodiversity.

## Introduction

Tropical rainforests are indispensable components of Earth’s systems, playing critical roles in maintaining global biogeochemical cycles and supporting human well-being through essential services such as food provisioning, water quality regulation, and global climate regulation (*1*). Tropical rainforests also harbour a tremendous and unique reservoir of biodiversity, a vast part being largely unexplored (*2*, *3*). Within these ecosystems, the forest canopy is a particularly dynamic and significant zone, representing the active interface between the terrestrial biosphere and the atmosphere, and a recognized hotspot of biodiversity (*4*). However, canopies’ inaccessibility has historically limited their scientific exploration, leaving substantial knowledge gaps. With the increasing pressures tropical rainforests face, including deforestation, biological invasions, and climate change (*5–7*), improving methods to monitor rainforests biodiversity, and in particular that of their canopies is now a pressing societal demand.

In recent years, the amplification and sequencing of taxonomically-informative DNA fragments from environmental samples (*8–10*), i.e. eDNA metabarcoding, has revolutionised biomonitoring across all domains of life. Environmental matrices such as soil (*9*, *11*), invertebrates bulk (*12*, *13*), marine and freshwater (*14–16*), or even air (*17–19*), are being commonly sampled environmental matrices for such purposes. However, these environmental matrices present several caveats for the sampling of terrestrial aboveground biodiversity (*20*). Specifically, the DNA found in soil samples is dominated by that of belowground organisms and overall transport poorly, making the probability of detecting aboveground organisms very low (*9*, *20*). The ingested DNA (i.e. iDNA) contained in hematophagous and/or phytophagous arthropod bulks can effectively sample vertebrates or plant communities (*12*, *13*) but this approach is destructive and hence not desirable. Filtering of air (*17–19*) or of water from local streams or lakes (*14–16*) is less invasive, but present several uncertainties related to the spatio-temporal window of the signal retrieved (*21*, *22*).

Organisms inhabiting the canopy naturally release DNA through tissue renewal and excretion, releasing cellular debris, urine, feces, and other materials that can adsorb onto the vegetation surfaces, creating a valuable eDNA source for biomonitoring. As such, analysis of eDNA from leaf swabs has been shown as a promising approach to detect terrestrial vertebrates (*23*). However, translating this approach to large-scale biodiversity assessments in tropical rainforests necessitates a considerable sampling effort, demanding alternative sampling protocols of this material. The sampling of canopy throughfall water represents such an alternative, as rainwash water transports and concentrates the DNA adsorbed on canopy leaves and branches. Accordingly, both experimental and *in situ* studies have demonstrated that it was possible to detect canopy arthropod species directly from rainwash water collected directly below the canopy (*24*, *25*). Rainwash waters therefore appear as extremely relevant for implementing large-scale eDNA-based monitoring of tropical forest canopies, these ecosystems being further characterised by an equatorial rainfall regime, with abundant rainfall throughout the year, albeit seasonal drier periods.

Here, we assessed the potential of the DNA contained in rainwash water collected under the canopy for the monitoring of plants, vertebrates and insects diversity in tropical rainforests (Figure 1). To this end, we conducted two sampling campaigns in two contrasted amazonian forests: one old growth forest, and a rubber (*Hevea brasiliensis*, Euphorbiaceae) and rosewood (*Aniba rosaeodora*, Lauraceae) tree plantation (Figure 1a). To collect canopy rainwash water, we repurposed umbrellas (Figure S1) and equipped them with four types of passive eDNA samplers (Figure 1b). The first campaign, hereafter referred to as the “temporal experiment”, aimed at assessing the temporal dynamics of rainwash eDNA in terms of (i) diversity/composition in Molecular Operational Taxonomic Units (here after referred to as MOTUs) through DNA metabarcoding (Table S1), and of (ii) eDNA persistence through quantification of an exogenous DNA pulverised on canopy leaves at the start of the experiment, using both digital PCR (dPCR) and DNA metabarcoding. This exogenous DNA was obtained from carrot, this species, *Daucus carota*, belonging to Apiaceae, a family absent from the studied area. The second campaign, hereafter referred to as the “spatial experiment”, aimed at assessing the spatial heterogeneity of the signal and sampling coverage needed to maximise the representativeness of the plot-scale diversity retrieved by rainwash eDNA. To this end, we collected rainwash eDNA every 20 meters in each 1ha plot using the most effective eDNA passive sampler tested during the first sampling campaign.

**Figure 1:**
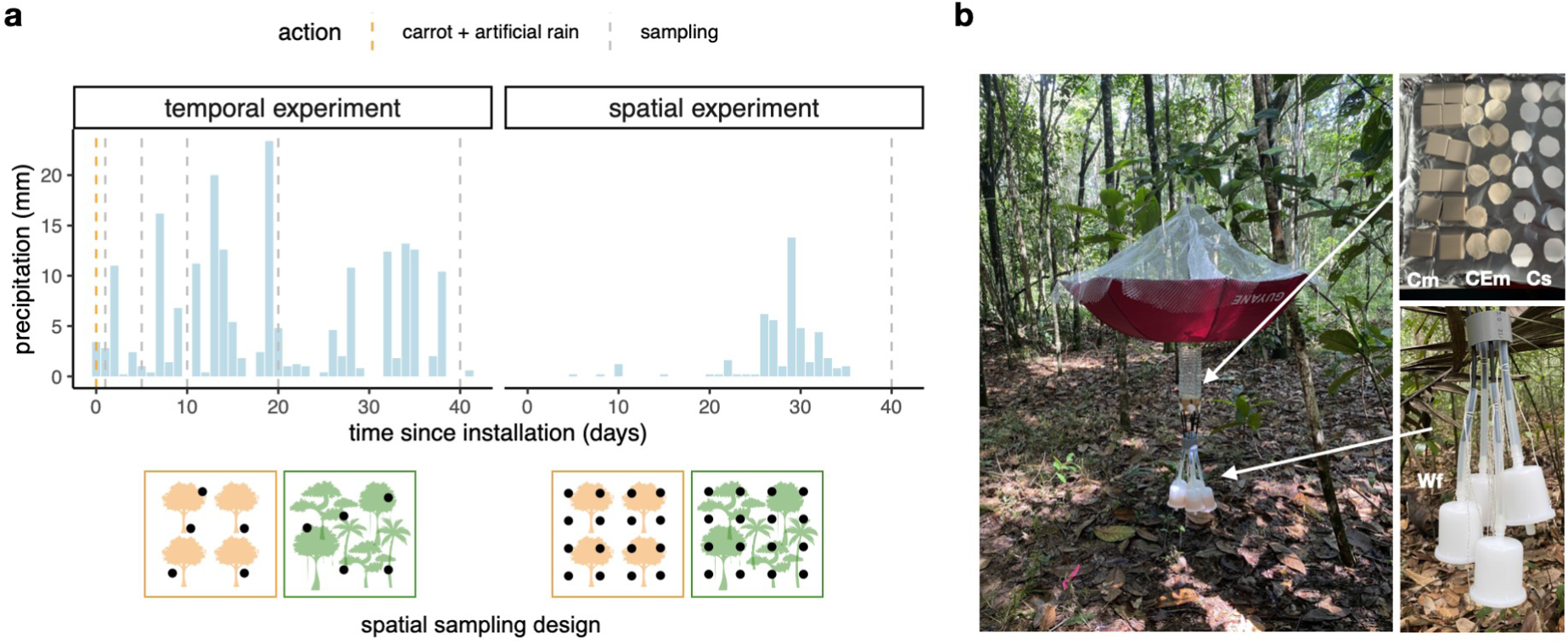
Overview of the study experimental designs. **a)** Description of the temporal and spatial experiments in terms of precipitation received throughout the sampling periods, initial setting (i.e. addition of carrot spike-in or not), as well as the spatial and temporal distribution of the sampling points. Orange plots: tree plantation, green plots: old-growth forest. **b)** Pictures of a canopy rainwash sampler (see also Figure S1) and of passive eDNA samplers (Cm: ceramic mosaics, CEm: Cellulose Ester membrane Cs: cotton strips, Wf: Waterra filters.

## Results and Discussion

### Rainwash eDNA captures canopy diversity

We first asked what was the overall taxonomic composition of the signal yielded by rainwash eDNA. Considering all samples from the temporal experiment, we were able to retrieve a total of 562 taxa (i.e. identified taxa - to any rank from phylum to species - regardless of the sequence similarity against its closer match in Genbank). Of these, the plant and vertebrate taxa identified to a fine taxonomic level are known to be present in South America (Figure 2, Table S2). For plants, we detected a total of 170 taxa, encompassing 50 orders and 56 species. These figures are low for a tropical forest, and can be due to the overall low resolution of the plant DNA marker used here, a portion of the *rbc*L gene. Thirteen of these taxa corresponded to mosses and liverworts, 14 to ferns including the arborescent ferns family, 3 to gymnosperms, and 16 to green algae. The remaining taxa corresponded to flowering plant families known to occur in the study area, such as Lecythidaceae, Melastomataceae, Fabaceae, Chrysobalanaceae, Vochysiaceae, Euphorbiaceae, Annonacae and, in much lower numbers, angiosperm epiphytes such as Bromeliaceae or Orchidaceae.

**Figure 2:**
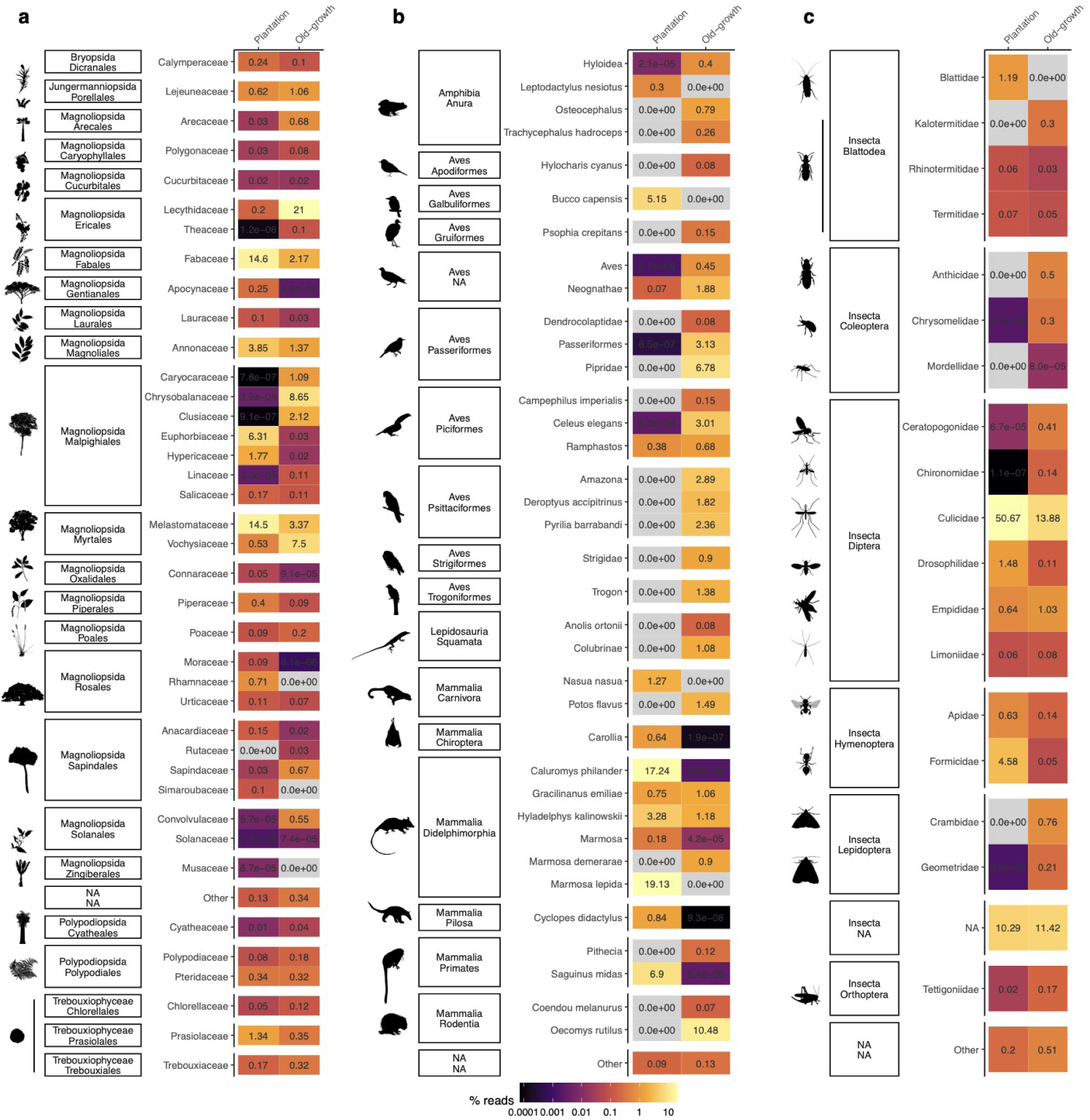
Taxonomic composition of canopy rainwash eDNA. In the tree plantation and the old-growth forest plots for **a**) plants, **b**) vertebrates and **c**) insects in the temporal experiment. Only taxa with a relative abundance (in terms of % reads) > 0.5% in the total dataset are shown (summarized at the family levels for plants and insects. The percentage of reads reported here corresponds to sequencing reads summed across all samples of the same condition (n = 25 per forest type, i.e. 5 spatial and 5 temporal sampling points). Grey cells correspond to null values, black ones to very small and yellow ones to high values.

Regarding vertebrates, after careful curation of common contaminants (human, beef and pig) and remaining tag-jumps (*26*, *27*) originating from positive controls, we retrieved a total of 72 taxa (spanning 21 vertebrate orders). These included 11 amphibian taxa, including the frog *Leptodactylus nesiotus*, known to occur in the area (*28*), 24 bird taxa (including e.g. parrots, toucans), four squamate taxa, and 32 mammal taxa. Most mammals corresponded to opossums and rodents, but also comprised two genera of neotropical monkeys (i.e. *Saguinus* and *Pithecia*), one genera of sloth (*Choloepus*) and of anteater (*Cyclopes*), two genera of bats (*Carollia* and *Rhinophylla*), and two genera of procyonids (i.e. *Potos* and *Nasua*). All taxa corresponded to species strictly or occasionally arboreal (*29*). We also detected two fish taxa. Fishes have also been detected on leaf swabs (*23*) but since it is unclear whether these corresponded to sporadic contamination or to the digestion product of another vertebrate, we excluded them from the analysis. Similarly, we detected sporadically the domestic cat. While it may correspond to true presences - the two plots being close from populated areas, we adopted a conservative filtration procedure and excluded these sequences from the analysis (see Methods).

For insects, we retrieved a total of 313 taxa, representing 20 orders. We also found 5 non-insect arthropod taxa (i.e. 2 spiders, 2 springtails, and one isopod). The most represented insect orders (in terms of diversity of taxa) were Coleoptera, Diptera (including several species of fruit- and sandflies, mosquitoes), Hymenoptera and Lepidoptera, which are also the most diverse orders in these ecosystems. Other insects such as cockroaches, termites, true-bugs, grasshoppers or dragonflies were also identified. Some of the taxa identified at a fine taxonomic are not usually found in the neotropical realm. However, the sequence identity scores with the closest matches in Genbank were overall quite low (i.e. below 90%) for insect taxa identified at either fine or coarse taxonomic resolution. This finding exemplifies the large incompleteness of the reference database and the resulting difficulty in achieving accurate taxonomic assignments, particularly for understudied tropical insect groups, and could explain at least partly why the number of retrieved arthropod taxa here was low compared to expectations for such environments (*30*).

We also investigated the relevance of the approach from a conservation point of view. Few taxa were in the vulnerable to critically endangered status, but a careful consideration of their identity and of their distribution range reveals that they do not occur in French Guiana, and most likely correspond to the mis-assignment of sister taxa occurring locally but that are not referenced in DNA databases, leading to an erroneous taxonomic assignment at the species level (i.e. the local species belonging to the genera *Campephilus, Dipteryx,* and *Swartzia* species were wrongly assigned to *Campephilus imperialis, Dipteryx alata* and *Swartzia bahiensis,* respectively). This illustrates how important it is to carefully verify the taxon list retrieved from eDNA surveys, as well as the need for pursuing efforts for populating DNA reference databases, particularly for tropical diversity, which remains largely under-represented (*20*, *31*). The rest were either found without IUCN status or were explicitly not evaluated, thus exemplifying the usefulness of rainwash eDNA to document the distribution of data deficient taxa, and its relevance for conservation efforts since over half of data deficient species are projected to face extinction (*5*).

Finally, we assessed the ecological relevance of the signal retrieved. We found the taxonomic diversity and composition in the two types of forest to be consistent with our expectations. First, the taxonomic diversity was 1.3 to 1.9 times higher in the old-growth forest plot depending of the taxonomic group, with 155 plant taxa (vs. 111 in the plantation), 61 vertebrate taxa (vs. 32) and 276 (vs. 153) insect taxa. The retrieved taxonomic composition also mirrored the contrasted disturbance status of the two forest plots (Figure 2). The rubber plantation plot yielded a higher proportion of reads assigned to family Euphorbiaceae (to which the rubber tree belongs), with its most representative MOTU corresponding to the rubber tree itself (*Hevea,* Table S2). We also detected greater abundance of termites belonging to the genus *Coptotermes*, to which the rubber termite belongs. In the plantation plot, we also detected a higher relative abundance of Lauraceae (to which rosewood belongs). We also found a higher proportion of reads assigned to Fabaceae, Melastomataceae, Hypericaceae, and Piperaceae, as well as to the frog *Leptodactylus nesiotus*. All these taxa are often associated with disturbed or more open habitats (*28*, *32*). For insects, we also found a higher relative abundance of mosquitoes, ants and cockroaches in the rubber plantation. Conversely, the old-growth forest plot harboured higher relative abundance of Lecythidaceae, Chrysobalanaceae, Vochysiaceae, and Caryocaraceae, as well as the frog *Trachycephalus hadroceps*. These taxa are typically found in old-growth forests (*32*, *33*). The old-growth forest also harboured more MOTUs and sequences assigned to phytophagous taxa such as Lepidoptera and Coleoptera.

### Optimising the sampling of rainwash eDNA

Making rainwash eDNA-based biomonitoring operational for routine analyses by environmental stakeholders requires developing sampling devices and strategies (i) that are cost effective and - if possible - low-tech for higher transferability, (ii) that limit the number of sampling campaigns while (iii) being able to generate biodiversity inventories that are spatially and temporally representative of the study area. This assessment is even more important for an effective implementation in tropical countries, where challenges of accessibility to sites are more important.

### Passive collection of eDNA

We developed a low-cost eDNA collection method where rainwash eDNA accumulates passively. This can be done either by filtration, as commonly done in waterbodies (*14–16*) but here filtration was not performed actively by pumping, but passively by gravity. Passive eDNA collection can also be done through passive fixation of eDNA molecules on different types of matrices (*34*, *35*) (Figure 1b). We assessed the efficiency of five passive eDNA samplers in fixing rainwash eDNA after 1 day of immersion in rainwash water. First, we compared the MOTU diversity retrieved by the different types of passive eDNA samplers, here estimated as Hill numbers set at *q* = 1, i.e. a number of equivalent species downweighting rare MOTUs. This index tends to the exponential of the Shannon index. We chose it because it is more robust for DNA metabarcoding data, due to its lower sensitivity to sequencing depth (Figure S2) or sampling effort, but also to other molecular low-abundance artefacts (*36–39*). Second, we compared the retrieved amount of carrot spike-in DNA across eDNA sampler types (Figure 1a), either in terms of copy number of a single-copy gene, *Dau c1*, quantified through digital PCR (dPCR), or based on the average number of Apiaceae sequencing reads retrieved in metabarcoding data. We considered these two measures here because while they correlated well (*40*) (Figure S3), they also have their own *pros* and *cons* (see Methods).

Filtration by gravity with Waterra filters captured higher MOTU diversity for plants and to a lesser extent for vertebrates compared to the other passive captors (Figure S4a, Table S3). A similar trend was observed for invertebrates, but was not significant. This collection method was also more effective in fixing the carrot spike-in DNA, followed by cotton strips after 1 day of immersion in rainwash water (Figure S4b, Table S3), but also for longer immersion periods (Figure S5). Hence, we recommend to use Waterra filters with filtration by gravity to maximise the sampling of rainwash eDNA. Fixation on cotton strips, which is more affordable, could be considered as an effective alternative collection method, but will require greater fixation surface and/or more passive sampler replicates per rainwash water samplers to reach a level of detection similar to passive filtration with Waterra filters.

### Temporal coverage of rainwash eDNA

Sampling tropical diversity effectively with rainwash eDNA also requires identifying the temporal sampling window that maximises the diversity retrieved due to an accumulation of rain events (Figure 1a), and of eDNA molecules coming from the canopy, but at the same time without losing too much information due to eDNA decay. Here, we evaluated this property on the Waterra filters obtained from the temporal experimental setup (Figure 1a) by considering three criteria: MOTU diversity dynamics, MOTU compositional temporal turnover, and spike-in DNA decay.

We analysed how MOTUs diversity of each target clade varied with the duration of the eDNA collection. For this specific analysis, we measured MOTU diversity at *q* = 0 (i.e. MOTU richness) on rarefied data because of the higher sensitivity of this index to subtle changes in species diversity and composition that would be caused by rare species (*38*), in our case of MOTUs becoming rarer due to eDNA decay. We fitted several functions modelling population dynamics (*41*) on the temporal trends of MOTU diversity and found that the Ricker function, which exhibits an optimum, was the best fit (Figure 3a). The fact that the temporal dynamics of MOTU diversity exhibits an optimum can be attributed to the reduced influxes of new eDNA molecules through time resulting from fewer and less intense rain events (Figure 1a), coupled with a stable eDNA degradation. Based on this model, we estimated the initial MOTU accumulation rate, which here ranged between 5 and 10 MOTUs per day after installation, as well as the time at which the diversity is maximal (Figure 3b, Table S4), here between 8 to 20 days after the installation of rainwash eDNA samplers.

**Figure 3:**
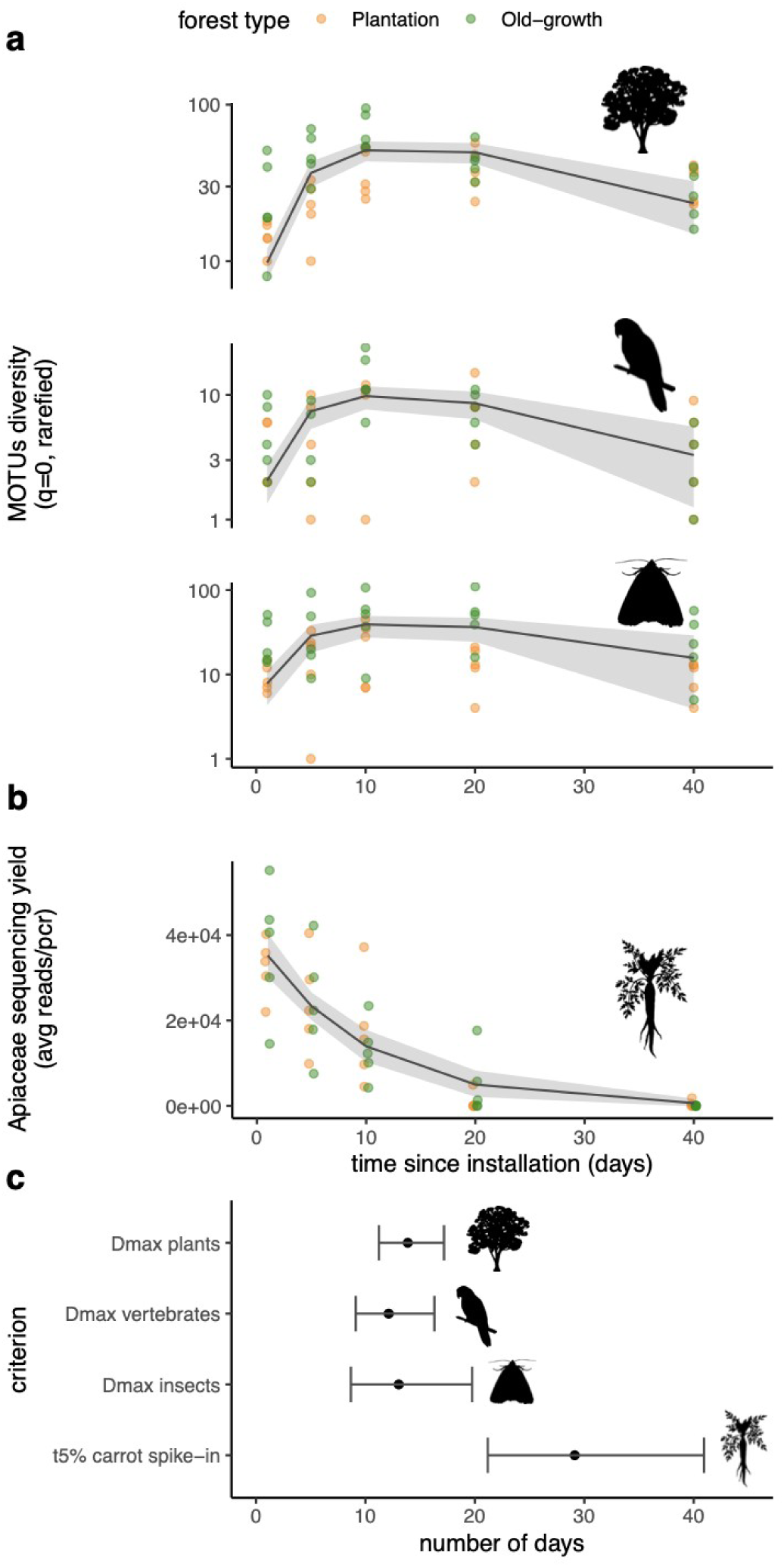
Dynamics and optimal temporal sampling window of rainwash eDNA. Dynamics of **a**) MOTU diversity (q=0, rarefied data) of plants, vertebrates or insects, **b**) amount of carrot spike-in DNA. Grey lines and grey transparent ribbons in (**a-b**) correspond to the models’ fits (see also Table S4) and 95% confidence interval. **c**) Temporal window of detection in terms of maximum diversity for each target clade or of 5% of spike-in initial concentration, with confidence intervals at 95%.

We also identified a time-decay pattern, i.e. a decrease of MOTUs compositional similarity as the duration between temporal sampling points increases (Figure S6). This supports the idea that the observed diversity dynamics of MOTUs is a balance between eDNA persistence in the matrix and influx of new molecules. However, this relationship was weaker for animals, and statistically non-significant for insects. We explain this by the fact that (i) several samples collected at close dates already exhibited very few to no shared animal MOTUs and that this phenomenon would be exacerbated using rarefied data, and (ii) all target clades exhibited a lower MOTUs diversity for the largest temporal intervals between sampling points (Figure 3). These properties may artificially inflate the weight of the MOTUs shared between samples in the similarity computation and of the similarity value itself, thus flattering the distance-decay pattern.

Finally, we estimated the decay rate of the carrot spike-in DNA using a negative exponential function (*42*), from which we estimated the time of decay to 5% of its initial quantity to about 29 days (see Methods, Figure 3a-b, Table S4).

We conclude that our passive sampling of rainwash eDNA can integrate relevant signal from 8 to 40 days depending on the initial amount of eDNA at the site, making it less punctual than one-shot samplings, as it is the case for the sampling of soil, water, and air or leaves swabs, which needs to be repeated. This property also makes the implementation of the protocol more feasible for routine analyses, either by reducing the need to frequent onsite visits, and/or by implementing monthly sampling for real-time monitoring, hence providing valuable insights into temporal trends and potential shifts in the studied area. These results are a first assessment of rainwash eDNA dynamics in real-life conditions, even though they miss processes involved in eDNA dynamics in general (*43*), such as the type from which the DNA detected originates (e.g. tissues, faeces, pollen). The frequency and intensity of rain events is an obvious important driver of these dynamics (see an example below), limiting the relevance of deploying the approach in areas that receive lower precipitation annually or during certain times of the year, such as seasonally dry tropical forests. We therefore believe that our estimate of rainwash eDNA decay should be assessed more systematically using a similar, simple approach with an exogenous DNA easy to obtain and monitor as we did here.

### Spatial distribution of rainwash eDNA

Finally, we assessed the spatial sampling effort required to sample diversity within one 1 ha plot. In our spatial experiment (Figure 1), rainwash eDNA was sampled every 20 metres in each plot during 40 days. Precipitations were strongly reduced during this experiment, resulting in an overall reduced detection power, especially for vertebrates. While we were able to detect several vertebrates taxa known to occur locally (e.g. *Saguinus*, *Oecomys*, or *Ramphastos*), the level of contamination was quite high. This illustrates the fact that infrequent or single rain events (artificial or not) capture insufficient amounts of eDNA for robust downstream analyses. Here, we adopted a conservative approach by restricting our spatial analysis to plant and invertebrate data, for which we built rarefaction curves of MOTU diversity (*sensu* Gotelli & Colwell (*44*), i.e. the average diversity observed when increasing the number of samples), this time with *q*=1 (i.e. exponential of the Shannon index) to obtain a more robust estimate of the plot-scale diversity (*36–39*) (Methods, Figure 4).

**Figure 4:**
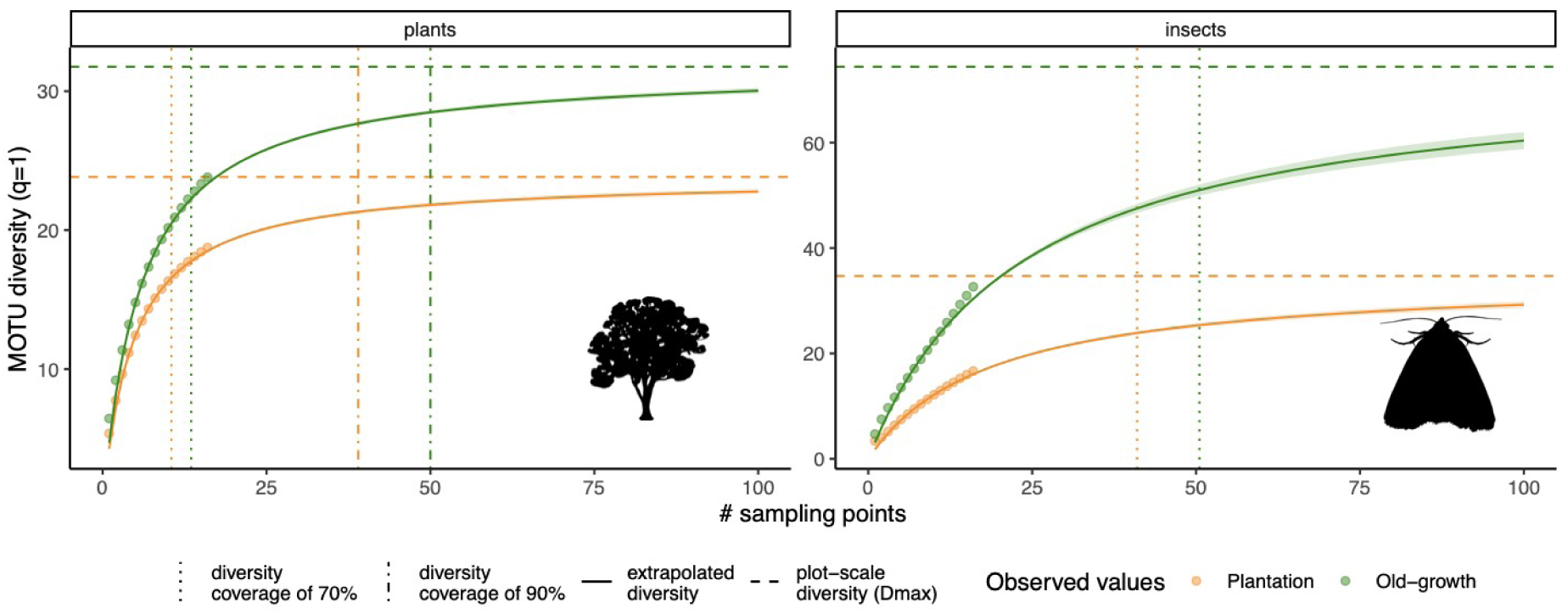
Accumulation of rainwash eDNA diversity with increasing sampling effort. For plants and insects. Points correspond to observed diversity values. The different line types represent parameters or curves derived from the Monod models’ fits (see also Table S5). Vertical lines correspond to the number of samples required to sample 70 and 90% of the plot-scale diversity. For insects, 90% of the diversity requires exceeds 100 samples (see Table S5). The transparent ribbons represent the confidence intervals at 95% of extrapolated diversity.

These rarefaction curves confirmed our previous observation (Figure 2) that the diversity recovered from rainwash eDNA is significantly higher in the old-growth forest for the targeted clades. This illustrates again the relevance of rainwash eDNA to detect differences of diversity across ecosystem types.

We then fitted species rarefaction models to the rarefaction curves (*45*) (Methods, Figure 4), from which we estimated the plot-scale diversity, the diversity covered by our current sampling effort (i.e. n=16 sampling points), as well as the number of sampling points required to cover 70% and 90% of the plot-scale diversity (Figure 4, Table S6). We found that these figures varied greatly across target clades, the plant diversity being relatively well covered (i.e. > 70%) with our current sampling effort. By contrast, we covered at best 46% of the insect plot-scale diversity detectable with our approach. While we do not exclude that these differences are partly due to the lower resolution of our markers (*9*), this result remain consistent with the tremendous diversity of insects usually described in tropical forests and at comparable spatial scales in comparison to that of plants (*30*). Reaching 90% of the plot-scale diversity would require at least 40 samples for plants, and much more for invertebrates (Figure 4, Table S6).

This suggests that the signal retrieved from rainwash eDNA, while acceptable to describe the diversity of non rare MOTUs at the plot scale, remains local, similarly to what is reported for soil eDNA (*20*) or from temperate rainwash eDNA (*25*). This conclusion is confirmed by the overall low to null MOTU compositional similarity between sampling points as well as the absence of a clear distance-decay pattern (Figure S7).

### Advantages of rainwash eDNA to monitor tropical rainforests

Our findings demonstrate that rainwash eDNA is a promising material for the monitoring of biodiversity in the tropical forest canopy by overcoming the challenges of working in a spatially heterogeneous environment both horizontally and vertically (*46–48*), and where eDNA is not transported as easily as in aquatic ecosystems. Our results show that to sample aboveground biodiversity, rainwash eDNA provides an effective non-destructive alternative to iDNA retrieved from arthropod bulks that has been also used for the same purpose(*12*, *13*). They also show that it offers a better representation of terrestrial above-ground biodiversity than soil eDNA, the latter being dominated by the eDNA of plants and soil organisms (*9*, *20*). Sampling the two matrices could hence provide a holistic view of both above- and belowground diversity.

Water eDNA from streams or lakes have been shown to provide useful information on terrestrial diversity (*14–16*). However, this material can only be collected in the presence of water bodies. In addition, water eDNA is transported across highly variable distances depending on a multitude of factors (e.g. particle size distribution, water body hydrogeomorphological properties, eDNA decay rate, etc.) (*22*). The water eDNA signal is also temporally variable, not only due to the demographic changes in the target biological community *per se*, but also because the fluctuation in frequency and intensity of rainfall events determines terrestrial material transport into water bodies (*49*). While also subjected to fluctuations in rainfall events, rainwash eDNA exhibits a localised spatial distribution because it is sampled closer to its source community. In addition, our study demonstrates that the community composition is detectable for at least two weeks. This suggests that the spatial dynamics of rainwash eDNA is easier to interpret than stream/lake water eDNA, and could offer a more localised and reliable approach for terrestrial biodiversity monitoring.

Air eDNA has also shown promise to describe terrestrial biodiversity, in particular that of vertebrates. However, current proof of concepts have been mostly conducted in zoos or similar systems with unvegetated soils - leading to high amounts of particles in the air, and/or harbouring a low diversity, and yet a high density of animals(*18*, *19*, *23*, *50*). These conditions facilitate animal detection and are not typical of tropical forests, where both animal and air particle density are expected to be much lower. In contrast, our assessment of rainwash eDNA has been done in natural conditions. In addition, air eDNA is likely to depend on the same spatio-temporal uncertainties as of stream/lake water eDNA (e.g. winds, rainfall events, etc) (*21*). Further experiments in natural conditions will be necessary to assess the advantages and/or complementarity of the two environmental matrices.

Apart from the limitations linked to molecular biases and contaminations that are common to all eDNA-based assessments (*9*, *51*), an important limit of our approach is the dependance to rainfall events. Given the persistence of rainwash eDNA found in moist tropical forests, this problem appears negligible for its appropriate sampling. However, it makes the approach less relevant for forests receiving less and irregular precipitation, such as tropical dry forests, cerrado forests or caatinga forests, where our approach would be relevant only during the wet season. On the other hand, we expect this approach to be very promising in temperate forests, where diversity is lower and where DNA will likely persist longer through time.

In conclusion, our study demonstrates the potential of rainwash eDNA as a powerful tool for monitoring the diversity of aboveground tropical forests’ plants and animals, which remains difficult to study with traditional and other eDNA-based methods. Our work establishes a robust methodological framework for an effective sampling of rainwash eDNA, and opens the door for further refinements for both community-level census and detection of patrimonial or human/environmental health importance. By enabling large-scale and non-invasive monitoring, this method can inform or assess tropical forests’ conservation strategies, and contribute to the protection of these irreplaceable ecosystems.

## Methods

### Study area

The study was conducted in the municipality of Sinnamary, which features two types of forest: an old-growth forest and a tree plantation. The equatorial climate in the studied area is characterised by two climatic periods: a long rainy period from mid-December to mid-August often interrupted by a short drier period between March and April and a longer dry season for the rest of the year. In this area, the annual rainfall is 3,041 mm with mean air temperature of 26°C (±1-1.5°C across the year). The 1 ha plot (plot P16) of old-growth forest is located in the Paracou Research Station (*52*) (lat: 5°18′N, lon: 52°53′W, https://paracou.cirad.fr/website). The Paracou research station managed by the French Agricultural Research Centre for International Development (CIRAD) comprises 5,000 ha of tropical rainforest in a hilly *terra firme* region. This site is a long-term research station where plant communities are well characterised and precipitation is monitored daily (Figure 1a).

The 1 ha plot of tree plantation is located in a former rubber tree (*Hevea brasiliensis*, Euphorbiaceae) plantation also managed by CIRAD, where rosewood (*Aniba rosaeodora*, Lauraceae) was more recently introduced and of which the exact current floristic composition is unknown. It is located 5 km away from the Paracou research station (lat: 5°17’N, lon: 52°54’W).

### eDNA sampling

#### Temporal experiment

To collect rainwash eDNA, we repurposed conventional umbrellas and plastic bottles into a low-cost rainwash water sampler with integrated passive eDNA sampler holders (instructions described in Figure S1). We equipped these samplers with four types of eDNA passive captors. First, we used Waterra eDNA filter (600 cm^2^ polyethersulfone filter, pore size of ⌀ 0.45µm, Waterra USA Inc.), which are commonly used for eDNA sampling in aquatic ecosystems (*53*) and that we here consider as a reference. We plugged the Waterra filters at the bottom of the water tank of the rainwash water sampler with silicon tubes to passively collect eDNA through gravity filtration (*54*) (Figure 1b). We also tested 3 alternative, cost-effective passive captors based on eDNA fixation. First, we considered Mixed Cellulose Ester membrane filters (⌀ 47 mm, pore size of 0.45µm, Merck KGaA, Darmstadt, Germany), which have recently proved effective for studying aquatic biodiversity with eDNA (*34*). We also considered ceramic mosaics of 25×25 mm, as ceramic has DNA-binding properties, even over long time periods (*55*, *56*). Finally, we considered cotton strips of 25×25mm, which also have the potential to passively sample aquatic eDNA (*57*). Ceramics and cotton strips have also long been used to measure biofilm colonisation/productivity rates (*58*) and decomposition (*59*, *60*), respectively. As biofilms also have eDNA-fixing properties (*35*), this property may increase the efficiency of these passive eDNA samplers for our purpose. All passive eDNA samplers were placed into the rainwash collector water tank.

A total of 5 rainwash water samplers were installed in each 1 ha forest plot at randomly selected locations away from the plot edges. We assessed the efficiency of the passive captors and quantified eDNA persistence through time by introducing an exogenous DNA just before the installation of the rainwash water samplers. We chose carrot (*Daucus carota*, *Apiaceae*) as a spike-in due to its widespread availability and because this plant family is absent from the study area and more generally from the Guiana Shield (except the aromatic plant *Eryngium foetidum*, which is not cultivated near the study sites (*61*)). To do so, we pulverised about 10 mL of carrot juice on canopy leaves at each sampling point with a conventional streamer. About 20 minutes after pulverisation, we installed below pulverised canopy leaves our rainwash water samplers, and then simulated an initial rain by pulverising around 40 mL of water (made of rainwater collected on site a few days before the start of the experiment (2:5) complemented with bottled springwater (3:5) to obtain a sufficient volume). eDNA passive captors were sampled after 1, 5, 10, 20 and 40 days for subsequent molecular analyses and placed in 50 mL sterile falcon tubes (Waterra filter excepted). We then added 50 mL of Longmire buffer (for 1L of buffer: 100mL 1M Tris, 100ml 1M EDTA, 2mL 5M NaCl and 798 mL of molecular biology grade H20) in each falcon tube and Waterra filter to store and preserve eDNA passive captors at room temperature (*62*) before molecular analyses. We thus obtained a total of 200 samples (i.e. 2 plots, 5 sampling points, 5 sampling dates, 4 passive eDNA samplers) for the temporal experiment.

#### Spatial experiment

In the same plots, we conducted another sampling campaign in mid February 2024 where we placed one rainwash water sampler every 20 metres, resulting in 16 points per plot. Following the results obtained in the temporal experiment, the water samplers were equipped with one Waterra filter each. The sampling devices were left at the field for 40 days (Figure 1). We thus obtained a total of 32 samples for the spatial experiment.

### Molecular analyses

#### DNA extractions

Each falcon tube containing a passive eDNA sampler immersed in Longmire buffer was shaken with a horizontal shaker at 40 Hz for 10 min solubilize the DNA, after what we removed the captor matrix. We also shaked the Waterra filters’ samples at 40 Hz for 10 min, after what we transferred the buffer into 50 mL falcon tubes. We then added 33 mL of absolute ethanol to all tubes and subsequently stored them at −20°C overnight. The following day, the tubes were removed from the freezer, left for 30 min at room temperature, and centrifuged at 8500 for 15 minutes at 15°C. The supernatant was then discarded, and 720µL of ATL buffer (QIAGEN; Hilden, Germany) was added to each pellet. The mixture was then transferred to 2 mL tubes, to which we added 20 µL of proteinase K (QIAGEN; Hilden, Germany), followed by an incubation at 56°C for 2 hours for digestion. NucleospinSoil® Macherey-Nagel kit was used, following the protocol from step 6 (adjust binding conditions) onward. The resulting DNA extracts were finally analysed using a Nanodrop ND-1000 (Thermo Fisher Scientific, Inc.) to determine the yield and purity of the DNA.

#### DNA metabarcoding

We characterised the community composition of plants, arthropods and vertebrates through eDNA metabarcoding, using the Sper03 marker (targeting a subregion of the rbcL gene, fwd 5’-CACCACAAACAGARACTAAARC-3’, rev 5’-TCCAYACRGTTGTCCATGTACC-3’) for plants, the Inse01 marker (*9*) (fwd 5’-RGACGAGAAGACCCTATARA-3’, rev: 5’-ACGCTGTTATCCCTAARGTA-3’) for insects, and the Vert01 marker (fwd 5’-TTAGATACCCCACTATGC-3’, rev 5’-TAGAACAGGCTCCTCTAG-3’) for vertebrates, together with a human-blocking primer: CTATGCTTAGCCCTAAACCTCAACAGTTAAATCAACAAAACTGCT-Spc3 (*63*). PCR reactions were performed in triplicate for each sample in 20 µL. Each reaction consisted of 10 µL of Amplitaq Gold 360 Master Mix (Applied Biosystems), 2 µL of a mix of fwd and rev primers (0.5 µM each final), 6 µL of Nuclease-free water, and 2 µL of eDNA diluted 10 times beforehand. For each PCR reaction, primers were labelled with unique combinations of 8-nt-long tags at the 5′ end of the forward and reverse primers (*64*) to distinguish PCR replicates after sequencing. PCR thermocycling conditions are indicated in Table S1. The final molecular experimental design thus included for each marker 600 PCR reactions corresponding to samples, 33 corresponding to extraction blank controls, and 30 to PCR blank controls for the temporal experiment. For the spatial experiment, it contained 96 PCR reactions corresponding to samples, 8 to extraction blank controls, and 3 to PCR blank controls. All PCRs of the same marker were pooled in equal volumes without standardising the PCR products concentration to allow a more quantitative comparison of the abundance of taxa sequencing reads across samples (*40*) (see also Figure S3). This pool was then used to build sequencing libraries using the Illumina Truseq Nano HT (Illumina), following the manufacturer protocol but without PCR enrichment of ligation products to minimise the formation of tag-jumps (*65*). Sequencing was conducted at the GetPlage sequencing platform (Toulouse, France) for the temporal experiment, and at Novogene (Cambridge, UK) for the spatial experiment, with the characteristics indicated in Table S1.

### Bioinformatics and data preparation

Primary bioinformatics/curation procedures, i.e. reads pairing, demultiplexing, dereplication, exclusion of obvious artefacts and singletons, and clustering of sequences to form Molecular Operational Taxonomic Units (MOTUs) were made with the `OBITools v1.2.11` (https://anaconda.org/bioconda/obitools, (*66*)) and the `sumaclust v1.0.31` clustering algorithm (https://anaconda.org/bioconda/sumaclust, (*67*)), using a snakemake workflow (https://github.com/AnneSoBen/obitools_workflow). The taxonomic assignment was done with the `ecotag` program (*66*) with the reference databases indicated in Table S1. These reference databases were built with the `ecoPCR` program (*68*) from Genbank. Next, we conducted a secondary curation procedure on sequencing data with the `metabaR R` package (https://github.com/metabaRfactory/metabaR, (*69*)), in order to further curate the data from contaminants, tag-jumps or other degraded sequences. Parameters used in this process are summarised in Table S1. Associated codes and diagnostic plots/statistics are available (Key resource Table). Finally, for vertebrates and plant data, we also verified that the taxon names retrieved had occurrence records in South America using the R package `rgbif v3.8.0` (https://CRAN.R-project.org/package=rgbif, (*70*)) of the Global Biodiversity Information Facility (GBIF, www.gbif.org). We also manually verified the distribution of each MOTUs in the PCR plate design to ensure they did not result from tag-jumps from positive controls. In some cases, we further compared their sequences against Genbank (assessed in Aug and Oct 2024) using BLAST (*71*). An overview of the dataset size at the main steps of the data curation process is shown in Table S6.

At the end of the data cleaning procedure, we checked how MOTUs diversity varied with sequencing depth for all markers and samples (Figure S1). While rarefaction curves did not saturate when considering MOTU richness, they did so when weighting MOTUs by their relative abundances using the Hill number framework (*72*, *73*). The Hill numbers, which correspond to an equivalent number of taxa, is more appropriate to analyse diversity with DNA metabarcoding data when its *q* order is > 0 (*36*, *37*). We thus present here alpha and beta diversity analyses for Hill’s *q* = 1 in most cases, corresponding to the exponential of the Shannon index for alpha diversity, and the Horn dissimilarity index for beta diversity (*73*). We did so for all analyses, except for those related to temporal MOTU diversity and compositional dynamics, where accounting for rare MOTUs is important due to eDNA decay processes. For this specific analysis, we computed diversity indices at q = 0, corresponding to MOTUs richness for alpha diversity, and the Sorensen index for beta diversity, but only after data rarefaction at 1000 reads / sample. All Hill numbers were calculated with the `hillR R` package `v0.5.2` (https://CRAN.R-project.org/package=hillR, (*74*)). Finally, we retrieved the IUCN status - when available - for all the taxa found using the `R` package `gbif.range` (https://github.com/8Ginette8/gbif.range, (*75*)). Note that we obtained a reduced diversity per sample overall for the spatial experiment ((Figure S1, Table S6), most likely due to the rarity of rain events during the spatial experiment (Figure 1). We further found that the vertebrate data was excessively contaminated. Because the contaminations pertained after rerunning the PCRs from scratch while most PCR blank controls were negative, we concluded that our DNA extracts were compromised for analysing vertebrates, either due to much rarer traces of local DNA than in the temporal experiment, or due to higher amounts of contaminations, or both. Consequently, we excluded this dataset from downstream analyses.

### dPCR assays

We also quantified the amount of carrot DNA for each passive eDNA sampler obtained at each sampled spatial and temporal points through digital PCR (dPCR) on a gene specific to carrots. For the dPCR-based approach, we quantified the copy number of the carrot *Dau c1* gene using the QIAcuity Digital PCR System (Qiagen; Hilden, Germany). The single copy gene targeted encodes for an allergen protein and is now routinely traced through PCR. In this study, we used the primers Dauc1F (5’-CCAGAGCCATTCACTCGAGATC-3’) and Dauc1R (5’-ACTGTATCAACATCAAGGACAATGC-3’) (*76*), together with the the QIAcuity EG PCR kit 25mL and QIAcuity Nanoplate 26k 24-well kits (QIAGEN; Hilden, Germany).

Each dPCR reaction was performed in a final volume of 40 µL, consisting of 13.3µL of EvaGreen, 3.2 µL of the two primers mixed at 0.4 µM final concentration each, 13.5µL of water and 10 µL of template DNA. We also performed one positive control per plate, adjusting the above conditions for 1 µL of carrot juice DNA extract as template DNA. Thermocycling conditions were as follow: an initial denaturation step at 95°C for 2 min, followed by 50 cycles of denaturation at 95°C for 15 sec, primer annealing at 58°C for 15 sec, and elongation at 72°C for 15 sec, and a final elongation step at 72°C for 5 min. The results were obtained from the machine with default parameters. For a selection of samples (i.e. Waterra filters obtained at t1 and t20 only), we also conducted dPCRs with the plant Sper03 marker (see Key resource table) using the reaction mix as above except that we adjusted the annealing temperature at 50°C (Table S1)

### Statistical analyses

All analyses performed with R were done with the R version 4.4.1 (*77*) and the tidyverse package (*78*).

### Taxonomic composition overview

To gain a first overview of the relevance of the taxonomic composition obtained from rainwash eDNA, we summed all MOTUs counts across sampling temporal and spatial points for each forest type of experiment 1 data. We aggregated the data at the plot scale rather than considering each spatial point individually due to the punctual information retrieved in each sample (see below for a more formal analysis on this aspect). We also aggregated the counts of all MOTUs assigned to the same taxonomic name.

### Identifying the best passive eDNA sampler

We identified the best type of passive eDNA sampler based on capacity to quickly fix a high diversity of taxa and high amounts of carrot spike-in DNA (i.e. one day after passive eDNA samplers immersion in rainwash water).

For the analysis of diversity detected by each passive sampler type, we considered the diversity of MOTUs at *q=*1 as explained above. For the analysis of carrot spike-in DNA, we used both dPCR assay data (expressed as a number of copies per µL of DNA extract) and a subset of DNA metabarcoding MOTUs classified as *Apiaceae* (expressed as a number of *Apiaceae* sequencing reads per PCR reaction). Both were not normally distributed, which is why the downstream analysis mostly consisted of non-parametric tests. We found the two types of data to be well correlated (Spearman’s ⍴ = 0.56, p < 0.001, Figure S3a). A similar trend was also observed when comparing the gene copy number of our plant marker and the corresponding total Sper03 sequencing reads per PCR reaction for a subset of samples (Spearman’s ⍴ = 0.7, p < 0.001, Figure S3b). This adds to previous evidence that DNA metabarcoding number of reads can be quantitative to a certain extent (*40*) when PCR products are sequenced together on the same lane and were not subjected to concentration standardisation prior sequencing. Nevertheless, we present here the two data types as they do not have the same detection levels, the single-copy nuclear gene Dau c1 being less detectable than the multi-copy chloroplastic gene used for DNA metabarcoding of plants, but the latter being more likely to be distorted by different molecular biases.

We then assessed the difference of the diversity, which also did not comply the assumption of normality, or the amount of carrot spike-in DNA detected across passive sampler types with Kruskall-Wallis tests, followed by *post-hoc* Wilcoxon tests with FDR-adjusted p-values when the former identified significant differences. Based on the results obtained, we considered only the best passive eDNA passive sampler for downstream analyses. These statistical analyses were run with the `R` package `rstatix v0.7.2` (https://cran.r-project.org/package=rstatix, (*79*)).

### Rainwash eDNA temporal dynamics

To identify the optimal temporal window for passive sampling of rainwash eDNA, we assessed the dynamics (i) of both MOTUs diversity and composition as well as (ii) of the carrot spike-in DNA. We did not expect the two plots to differ for these properties and preliminary analyses using mixed models did not retain this random effect. We thus analysed the data without accounting for this factor.

For this specific analysis, we considered MOTU diversity at *q*=0 after data rarefaction (see bioinformatics and data preparation section) rather than at *q=*1. We did so because it gives more weight to rare species (*38*), and is thus more sensitive to detect MOTU accumulation through time by better accounting for MOTUs that would become rarer due to eDNA decay. We expected MOTU diversity to accumulate rapidly at the start of the experiment and then either (i) to accumulate more slowly if the carrying capacity of the filter is not reached at *t_n_* and/or if the decay rate of eDNA is lower than that of the influx of new ones, (ii) to reach a plateau if the carrying capacity of the filter is reached and/or if an equilibrium between the decay rate of eDNA and the influx of new ones, and finally (iii) a decrease of MOTU diversity when the decay rate of rainwash eDNA exceeds that of the influx of new eDNA molecules (e.g. due to reduced rain events). To this end, we fitted several non-linear functions (*41*), i.e. power (*D* = *at^b^*), monomolecular (*D = a(1-e^-bt^)*), or Ricker (*D = ate^-bt^*) on plants, vertebrates and insects MOTU diversity (*D*) patterns through time (*t*). We selected the models exhibiting the lowest AIC, here built with the Ricker function, from which we estimated the initial diversity accumulation rate (*a*) and time at which the diversity was maximal (*T_Dmax_ = 1/b*). Models predicted values as well as their associated standard deviations and confidence intervals were obtained with the `R` package `propagate v1.0.6` (https://CRAN.R-project.org/package=propagate, (*80*)).

We also analysed the MOTU compositional turnover at *q*=0 (see above) using a distance-decay relationship approach, as we expected samples collected at close sampling date to exhibit higher similarity than those collected at distant sampling dates, due to both eDNA persistence/decay and arrival of new eDNA molecules through time. The temporal distance-decay pattern was studied by considering only the pairwise comparisons involving the same spatial sampling point. It was assessed with a semi-log linear model, which is often used to model distance-decay patterns (*81*) and permutation tests to assess the slope significance with the `R` package `lmPerm` (https://CRAN.R-project.org/package=lmPerm, (*82*)). To circumvent the log transformation problem on null similarity values, we added the smallest non-null similarity value to all similarity values.

Finally we determined the persistence of rainwash eDNA by estimating the decay of the carrot spike-in DNA using a first-order decay model (i.e. a negative exponential function) of the form *C_t_ = C_0_e^-λt^*where *C_0_* is the initial eDNA concentration, *t* is the time in days and *λ* the DNA decay rate constant (per day), from which then derived the time of rainwash eDNA detectability as *t_0.05_ = log(0.05)/λ*, corresponding to the time it would take for the carrot spike-in DNA to fall below 5% of its initial concentration. While more complex decay models have been found to outperform the one considered here to estimate eDNA decay rate constants (*42*), our approach allows for cross condition comparisons more easily and appears to be biased only for very slow decay rates (e.g. at low temperature and in absence of light), which we do not expect to be the case here. We fitted the decay model only on *Apiaceae* average reads abundance per PCR reaction. We did not consider here the number of Dau c1 copies / µL DNA extract, as the detection was too limited to obtain reliable fits and because quantities obtained are expressed in a different unit than DNA metabarcoding data.

### Rainwash eDNA spatial coverage

To assess the optimal spatial sampling effort, we built rarefaction curves for each target clade from data obtained in the spatial experiment (Figure 1a), by averaging all diversity values obtained from all possible sample combinations at each sampling intensity (in number of sampling points per plot) and for each plot separately, as we expected to two plot to exhibit different diversity and spatial accumulation rates (*44*). Here, MOTU diversity was measured at *q*=1 instead of *q*=0 on rarefied data. The main reason for this choice lies in the robustness of the diversity measure at *q*=1 to differences in sequencing efforts and potential remaining errors (*36*, *37*), as indicated above. In addition, the rate of increase of the uncertainty associated with species addition at *q*=1 is reduced compared to *q*=0 (*38*, *39*). Finally, unlike the MOTU temporal dynamics analysed above, we did not seek to detect subtle changes in species abundances with increasing sampling effort.

We then fitted on MOTU accumulation curves two simple functions classically used to describe species-area/accumulation curves and exhibiting an upper asymptote in order to estimate the plot-scale diversity and from this, of our sampling coverage. These functions corresponded to the Monod (*DN = D_max_N/(k_N_+N)*) and Negative exponential (*D = D_max_ (1-e^-kN^)*) models (*45*), where *D* is the diversity, *D_max_* the upper asymptote, corresponding here to a theoretical plot-scale diversity detectable with our approach over the period of time considered in our study, *N* the number of samples, *k_N_* the “half velocity constant”, in our case the number of samples required to reach 50% of the plot-scale diversity, and *k* a MOTU accumulation rate. As for above, we selected the function with the lowest AIC, which corresponded to the Monod models in all cases. We used these models to predict the number of spatial sampling points required to sample 70% and 90% of the plot-scale diversity. Models’ predicted values and associated confidence intervals were obtained with the `R` package `propagate` (*80*) as above. Finally, we also evaluated the pairwise MOTU compositional similarities at *q*=1 between samples across or within each forest plot. We also assessed the relationship between MOTU compositional similarity and geographic distances graphically.

## Data and code availability

Raw and pre-processed amplicon and dPCR data as well as data quality reports and analyses scripts are all stored on data.InDoRES (will be provided upon manuscript acceptance). Any additional information required to reanalyse the data reported in this paper is available from the lead contact upon request.

## Acknowledgements

We thank the Paracou forest research station in French Guiana, managed and supported by CIRAD, UMR EcoFoG (https://paracou.cirad.fr), and financially supported by a French Investissement d’Avenir programme (Labex CEBA ANR-10-LABX-25-01). We also thank the Genotoul bioinformatics platform of Toulouse Midi-Pyrénées (Bioinfo Genotoul) for providing computing and storage resources. LZ is deeply indebted to Markus Gastauer and to the Fundação de Ciência, Tecnologia, Inovação e Desenvolvimento Sustentável GUAMÁ, from whom she benefited from the grant Bolsa Estímulo à Inovação (BEI) no 094-2023. The work was funded by the OFB (Office Français de la Biodiversité) and has benefitted from ‘Investissement d’Avenir’ grants managed by Agence Nationale de la Recherche (CEBA: ANR-10-LABX-25-01; TULIP: ANR-10-LABX-0041).

## Author contributions

All authors contributed to the conception of the research.

Fr.P. developed the rainwash water collectors.

A.-S.B., A.I., C.L., E.L., Fi.P., Fr.P., V.T, J.O and C.R.-H. performed the fieldwork.

Y.C., A.I., E.L., L.M., V.T., and U.S. performed the molecular work.

A.-S.B. performed the bioinformatics analyses with the help of L.Z. and J.R.

L.Z. performed the statistical analyses with the help of J. P.

A.I. and L.Z. obtained the funding and managed the project.

L.Z. wrote the first draft of the paper and all authors contributed to revisions.

## Declaration of interests

The authors declare no competing interests.

## Supplementary Material

### Supplementary Figures

**Figure S1:**
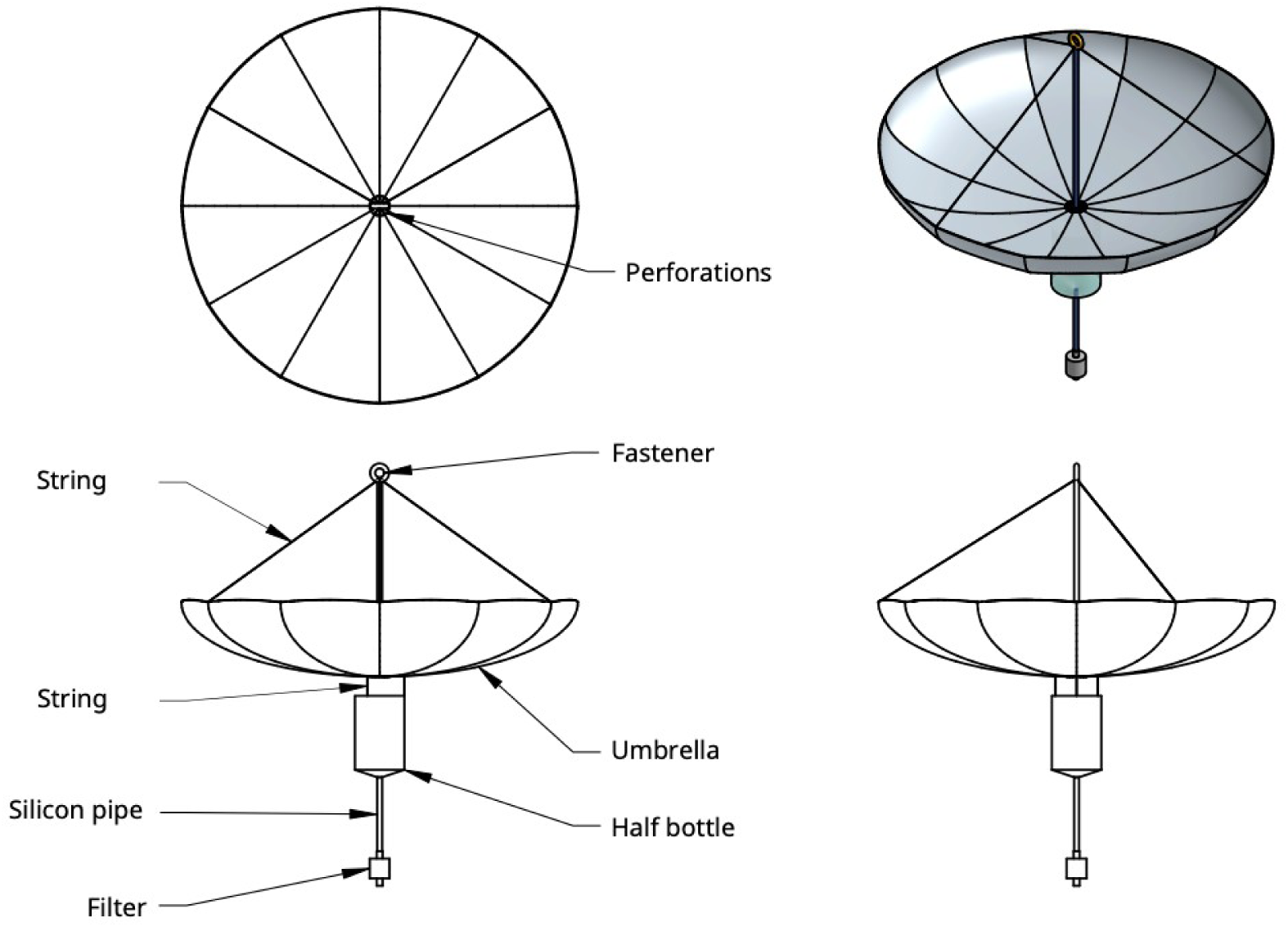
Construction diagram of rainwash water collectors from below (topleft), above (topright), a front view from different sides (bottom). The fastener is made of galvanized steel wire (1.5mm diameter), the strings are in 1.5mm nylon cord, the bottle corresponds to a classical 1.5L plastic bottle (here « Dilo » mineral water), and the filter corresponds to the Waterra filter.

**Figure S2:**
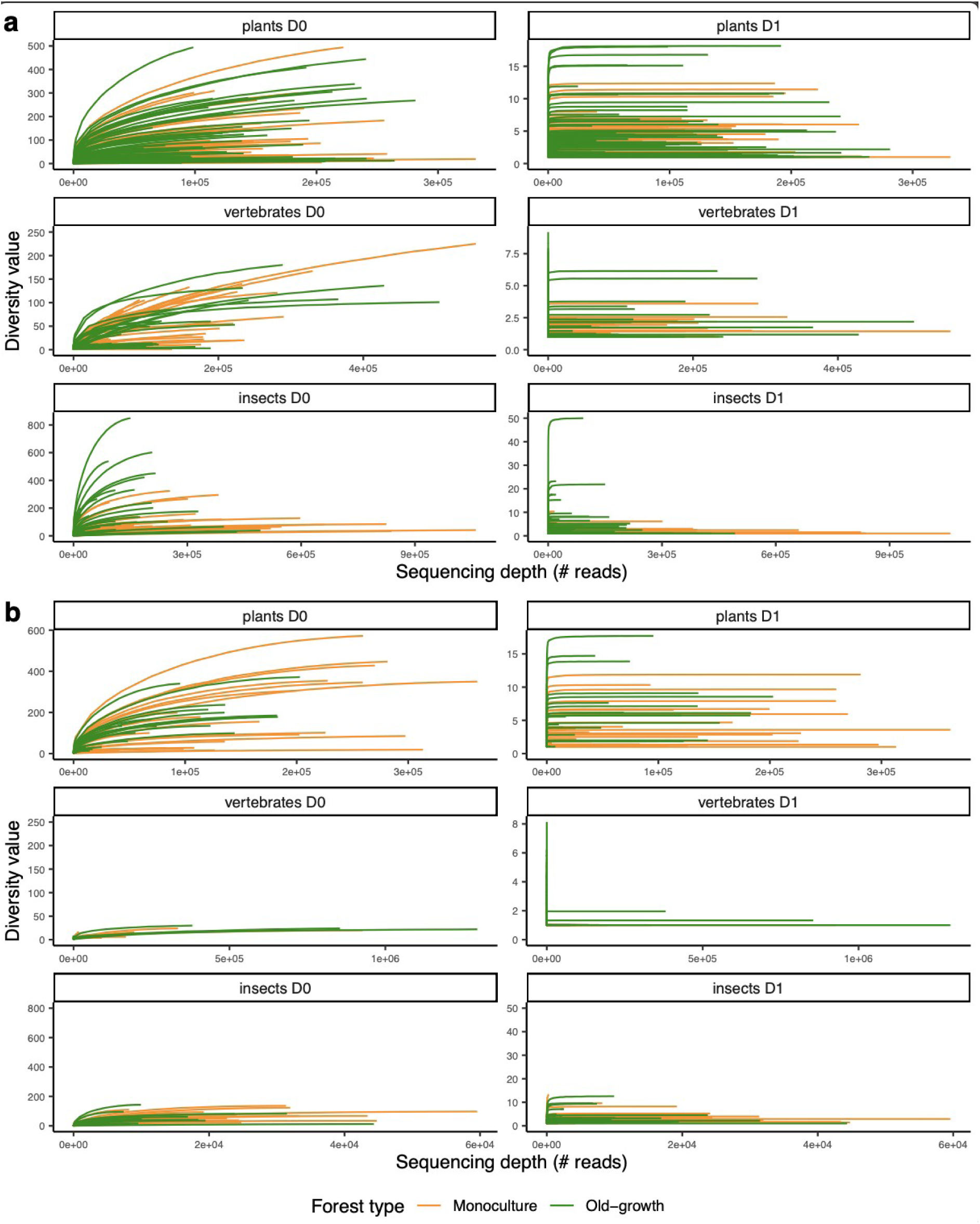
Rarefaction curves for plants, vertebrates, and insects metabarcoding data depicting the per-sample accumulation of MOTUs Hill’s diversity with increasing sequencing depth. D0 = observed MOTU richness (i.e. Hill’s *q*=0) and D1 ∼ exponential of the Shannon index (i.e. Hill’s *q*=1). **a)** Data from the temporal experiment, **b)** Data from the spatial experiment. “Monoculture” corresponds to the tree plantation.

**Figure S3:**
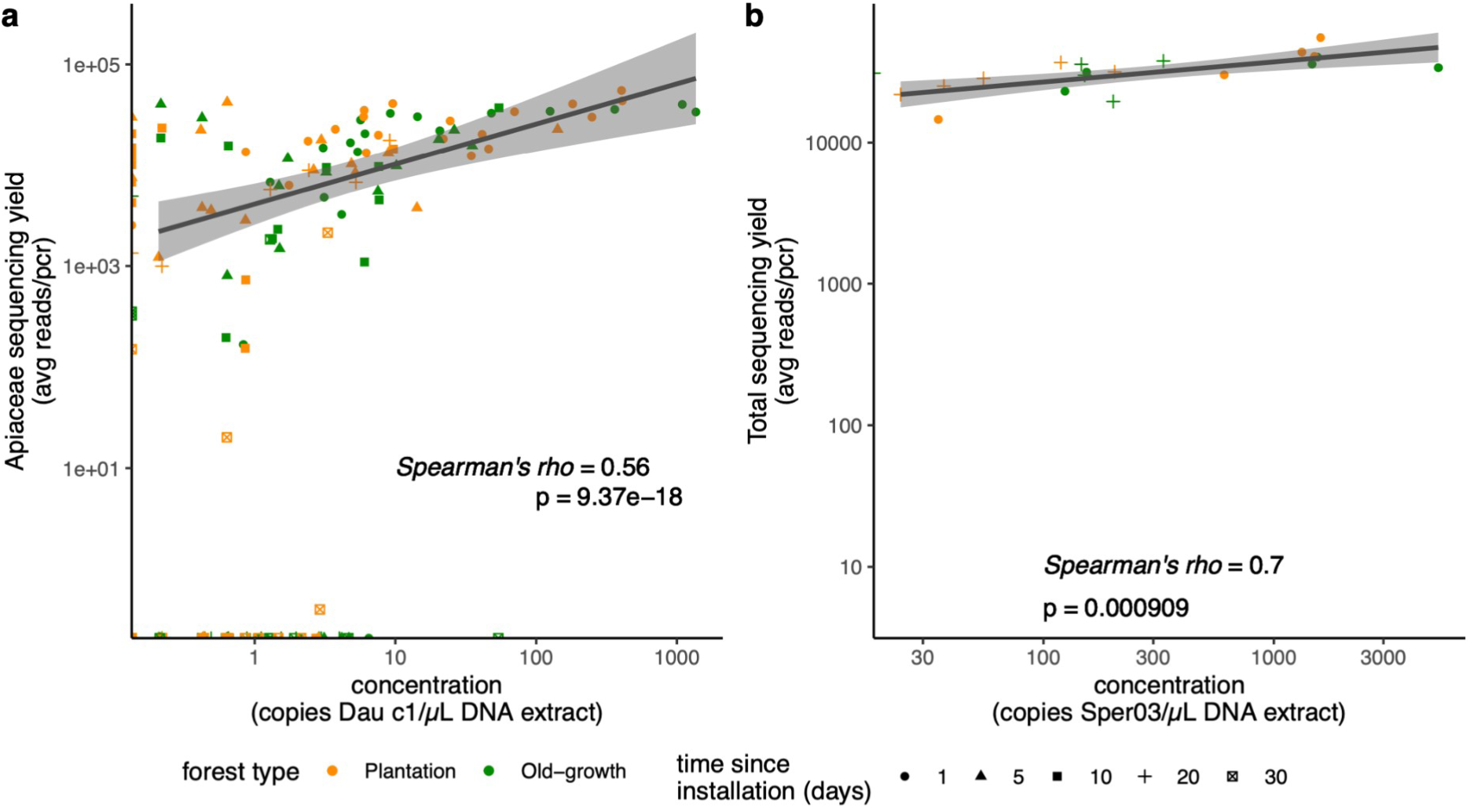
Correlation between gene copy numbers and sequencing yields per sample. For **a)** the *Dau c1* gene, as assessed through dPCR, and the average number of Apiaceae reads/pcr reaction (n=3 per sample) in DNA metabarcoding data for all samples analysed, and **b)** the Sper03 gene, as assessed through dPCR, and the average total number of reads/pcr reaction (n=3 per sample) in DNA metabarcoding data for Waterra samples. The Spearman correlation test statistics are also indicated.

**Figure S4:**
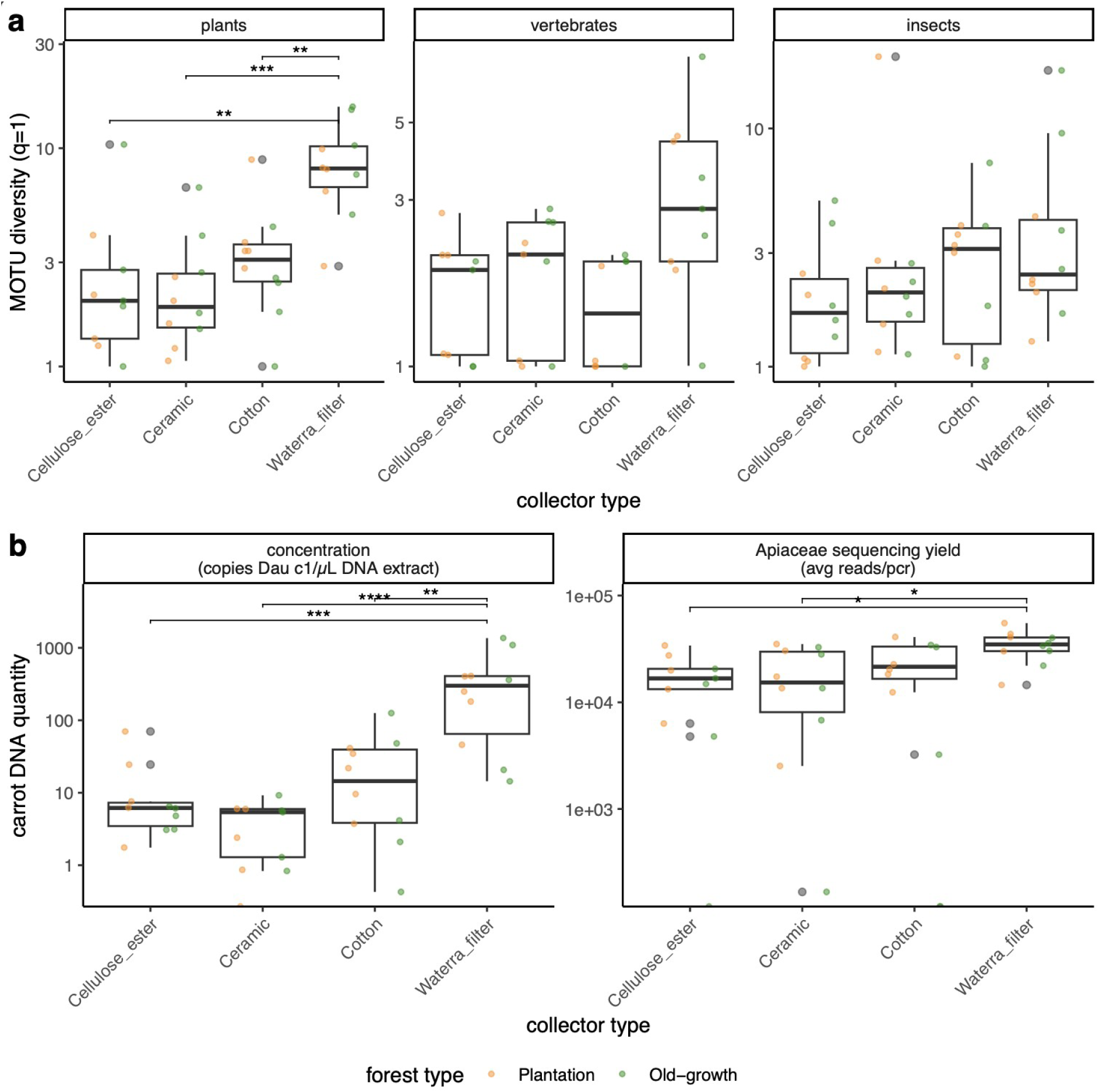
Efficiency of different passive eDNA samplers after 1 day of immersion in rainwash water in terms of fixation of: **(a)** MOTUs diversity (Hill’s *q*=1) and **(b)** carrot DNA expressed either as a number of *Dau c1* copies (i.e. dPCR data), or as a number of sequencing reads of *Apiaceae* (i.e. DNA metabarcoding data). Stars indicate the significant differences between passive eDNA sampler types identified in the Wilcoxon *post-hoc* tests (FDR-corrected p-values). Values from the tree plantation and the old-growth forest are shown with colours.

**Figure S5:**
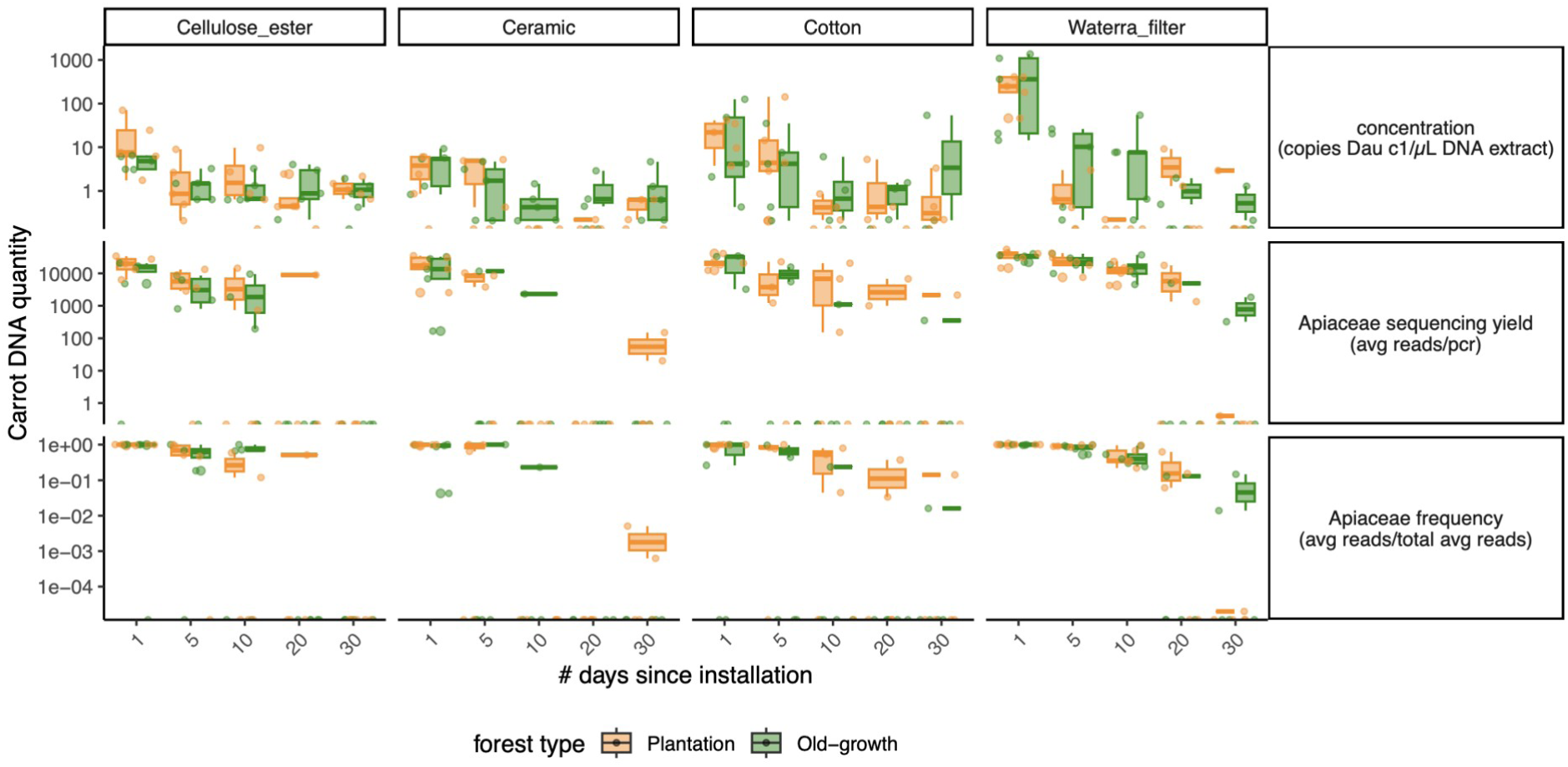
Overview of the amount of carrot spike-in DNA across types of passive eDNA samplers (columns) throughout the whole temporal experiment in terms of (rows) (i) number of copies of the nuclear single-copy gene Dau c1 per µL of DNA extract, (ii) average number of *Apiaceae* reads per PCR in DNA metabarcoding data, and (iii) average relative abundance of *Apiaceae* over the average total number of reads per PCR.

**Figure S6:**
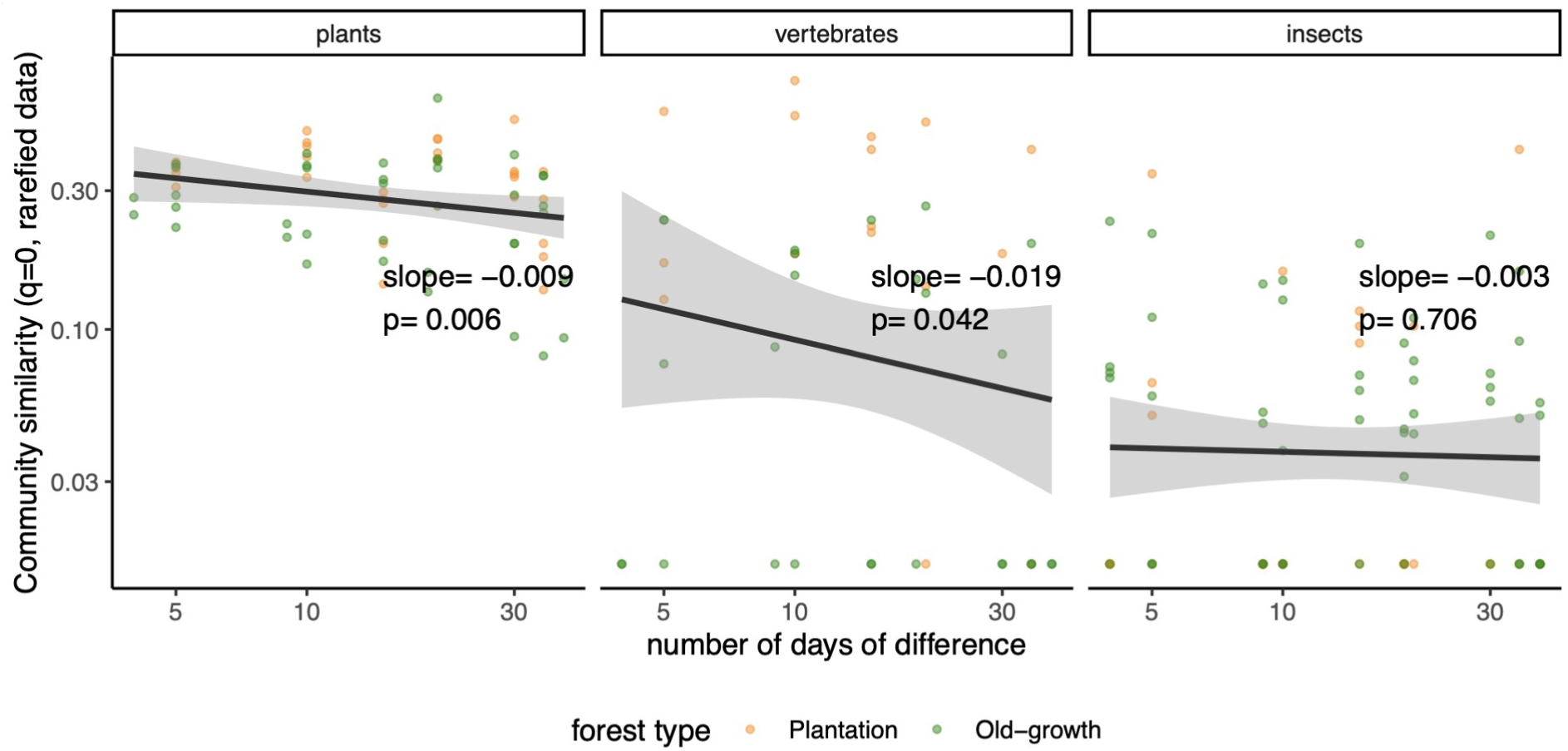
Temporal distance - similarity decay pattern of MOTU composition (as measured with the Hill number framework at *q=0* on rarefied data) across sampling dates for each spatial sampling point and for each target clade.

**Figure S7:**
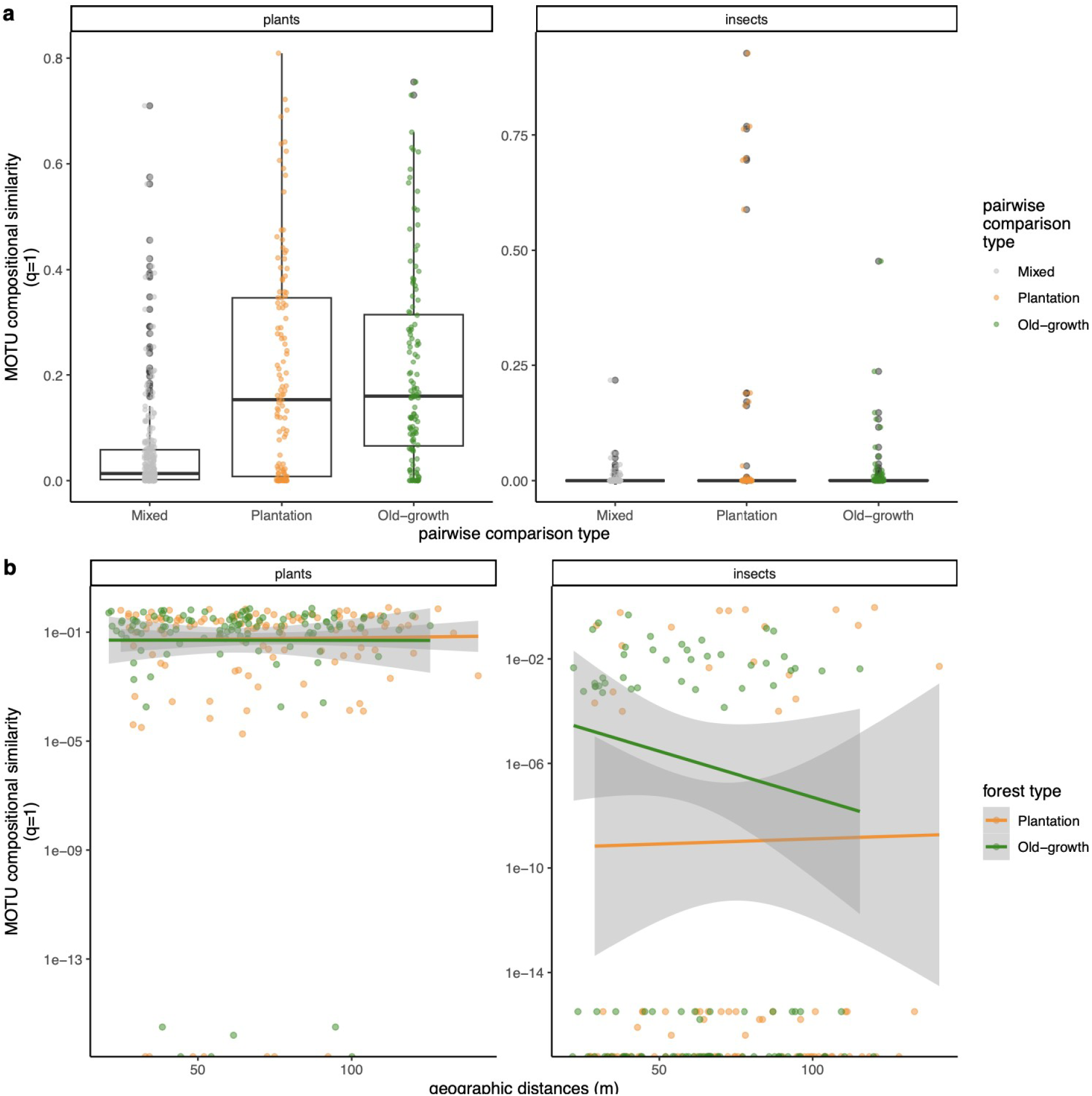
Spatial patterns of MOTU composition (as measured with the Hill number framework at *q*=1) across sampling plots and points for each target clade. **a)** MOTU compositional similarity between vs. within forest plots. **b)** Distance-decay patterns.

### Supplementary Tables

**Table S1:**
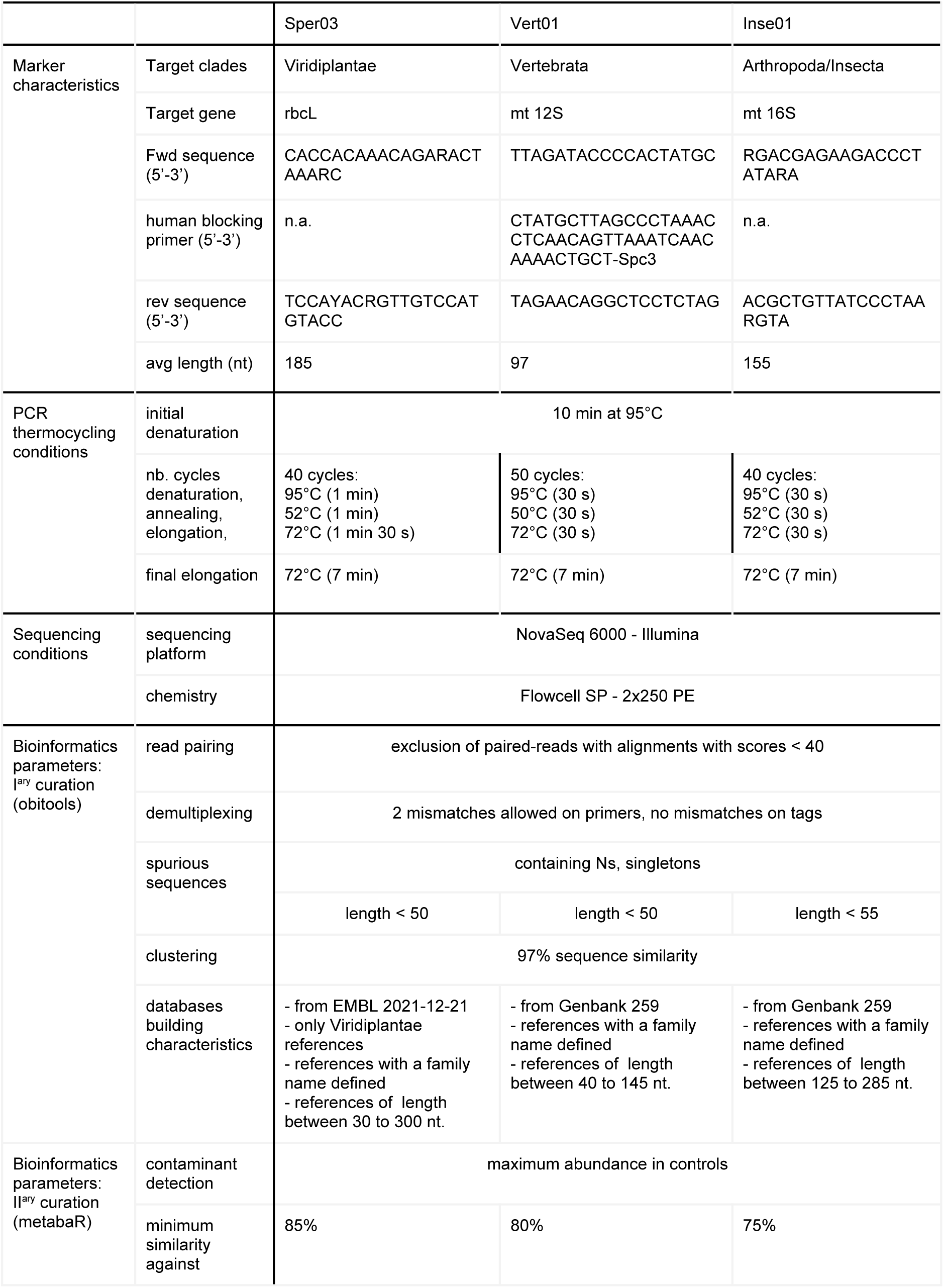
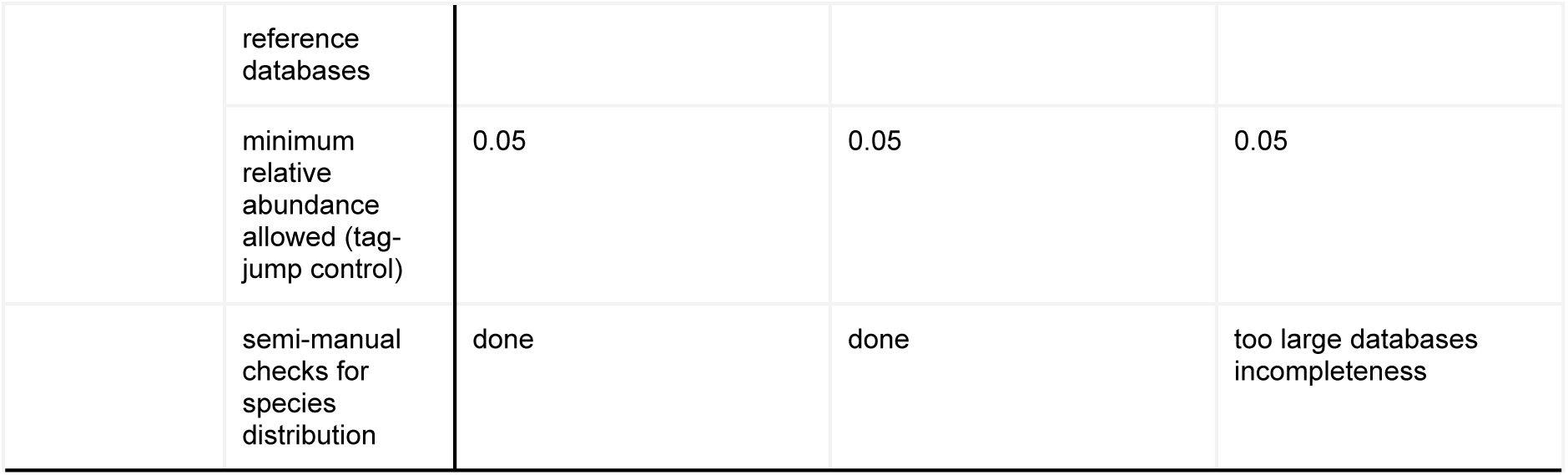
DNA markers and associated molecular and bioinformatics procedures and parameters used in this study. See Key resources Table for primers and software references.

**Table S2:** Full list of taxa retrieved in experiment 1 after data curation (=> table_metabar_taxalist_all.tsv)

**Table S3:** Kruskall and Wilcoxon test statistics comparing the amount of carrot spike-in DNA and MOTU diversity (q=1) for each target clade across passive eDNA sampler types. Abbreviations are as in Figure 1. Significant FDR-corrected p-values (p.adj) are in bold.

| Data type | Statistics |  |  |  |  |
| --- | --- | --- | --- | --- | --- |
| Plant MOTU diversity | Kruskal test<br>n = 30 | statistic | p-value |  |  |
|  |  | 13.2 | **7.53e-4** |  |  |
|  | Wilcoxon test<br>n = from 5 to 9 each | Passive sampler 1 (n) | Passive sampler 2 (n) | statistic W | p.adj |
|  |  | CEm | Cm | 47.5 | 0.870 |
|  |  | CEm | Cs | 30.5 | 0.304 |
|  |  | CEm | Wf | 9 | **0.004** |
|  |  | Cm | Cs | 32 | 0.285 |
|  |  | Cm | Wf | 4 | **7.8e-4** |
|  |  | Cs | Wf | 10 | **0.004** |
| Vertebrate MOTU diversity | Kruskal test<br>n = 35 | statistic | p-value |  |  |
|  |  | 8.54 | **0.0361** |  |  |
|  | Wilcoxon test<br>n = from 8 to 9 each | Passive sampler 1 (n) | Passive sampler 2 (n) | statistic W | p.adj |
|  |  | CEm | Cm | 30.5 | 0.479 |
|  |  | CEm | Cs | 42.5 | 0.56 |
|  |  | CEm | Wf | 16 | 0.101<br>(=0.034 non adjusted) |
|  |  | Cm | Cs | 52 | 0.198 |
|  |  | Cm | Wf | 22 | 0.198 |
|  |  | Cs | Wf | 11 | 0.101<br>(=0.018 non adjusted) |
| Insect MOTU diversity | Kruskal test<br>n = 40 | statistic | p-value |  |  |
|  |  | 3.46 | 0.327 |  |  |
| Concentration<br>(copies Dau c1/μL DNA extract) | Kruskal test<br>n = 40 | statistic | p-value |  |  |
|  |  | 20.9 | **0.000109** |  |  |
|  | Wilcoxon test<br>n = 10 each | Passive sampler 1 (n) | Passive sampler 2 (n) | statistic W | p.adj |
|  |  | CEm | Cm | 75 | 0.076 |
|  |  | CEm | Cs | 40 | 0.481 |
|  |  | CEm | Wf | 5 | **0.000618** |
|  |  | Cm | Cs | 25 | 0.076 |
|  |  | Cm | Wf | 0 | **0.0000648** |
|  |  | Cs | Wf | 12 | **0.006** |
| Apiaceae sequencing yield<br>(avg reads/pcr) | Kruskal test<br>n = 40 | statistic | p-value |  |  |
|  |  | 10.4 | **0.0151** |  |  |
|  | Wilcoxon test<br>n = 10 each | Passive sampler 1<br>(n) | Passive sampler 2<br>(n) | statistic W | p.adj |
|  |  | CEm | Cm | 44 | 0.880 |
|  |  | CEm | Cs | 45 | 0.880 |
|  |  | CEm | Wf | 11 | **0.013** |
|  |  | Cm | Cs | 49 | 0.970 |
|  |  | Cm | Wf | 16 | **0.027** |
|  |  | Cs | Wf | 21 | 0.062 |

**Table S4:** Coefficient estimates of MOTU diversity (q = 0 on rarefied data) accumulation and spike-in DNA decay (assessed here only with metabarcoding data) through time with their associated FDR-adjusted p-values. *a*: initial MOTU diversity accumulation rate, *TD_max_*: time at which MOTU diversity is maximal, **λ**: eDNA decay rate, *t_0.05_*: time of detectability at 5%, i.e. the time it would take for eDNA to fall below 5% of its initial quantity. n.a.: not applicable.

| Clade | Model type | Coefficient | Coefficient value | Confidence interval (95%) | p.adj |
| --- | --- | --- | --- | --- | --- |
| Plants | Ricker | $a$ | 9.86 | 7.50 - 12.76 | **3.86e-10** |
| | | $TD_{max}$ | 14.33 | 11.56 - 17.86 | **6.89e-13** |
| Vertebrates | Ricker | $a$ | 2.31 | 1.51 - 3.40 | **5.24e-06** |
| | | $TD_{max}$ | 11.74 | 8.60 - 16.19 | **4.21e-08** |
| Insects | Ricker | $a$ | 8.59 | 5.19 - 13.57 | **4.54e-05** |
| | | $TD_{max}$ | 13.14 | 8.94 - 19.46 | **1.08e-06** |
| Apiaceae sequencing yield (avg number of reads/pcr) | Negative exponential | $\lambda$ | 0.10 | 0.07 - 0.14 | **8.29e-07** |
| | | $t_{0.05}$ | 29.13 | 21.18 - 40.95 | n.a. |

**Table S5:** Coefficient estimates of the spatial accumulation of MOTU diversity (at *q*=1) with their associated FDR-adjusted p-values. *D_max_*: plot-scale estimated MOTU diversity, *N_D50_ (= k_N_), N_D70,_ N_D90_*: number of samples required to reach 50, 70 and 90% of the plot-scale diversity. *D_N16_*: Coverage (in %) of the plot-scale diversity with current sampling effort (n=16).

|  | Forest type | Tree plantation |  |  | Olg-growth |  |  |
| --- | --- | --- | --- | --- | --- | --- | --- |
| Clade | Coefficients | Estimate | CI (95%) | p.adj | Estimate | CI (95%) | p.adj |
| Plant | $D_{max}$ | 23.820 | 23.598 - 24.056 | 0 | 31.764 | 31.534 - 31.996 | 0 |
| | $N_{D50}(=k_N)$ | 4.602 | 4.482 - 4.723 | 0 | 5.771 | 5.671 - 5.872 | 0 |
| | $D_{N16}$ | 77.662 | 77.357 - 77.966 | n.a. | 73.492 | 73.288 - 73.696 | n.a. |
| | $N_{D70}$ | 10.5 | 10 - 11 | n.a. | 13.5 | 13 - 14 | n.a. |
| | $N_{D90}$ | 39 | 35 - 43 | n.a. | 50 | 45 - 55 | n.a. |
| Insects | $D_{max}$ | 34.675 | 33.699 - 35.711 | 0 | 74.399 | 71.729 - 77.282 | 0 |
| | $N_{D50}(=k_N)$ | 18.539 | 17.779 - 19.347 | 0 | 23.199 | 22.061 - 24.430 | 0 |
| | $D_{N16}$ | 46.324 | 46.022 - 46.624 | n.a. | 40.817 | 40.52 - 41.109 | n.a. |
| | $N_{D70}$ | 41 | 39 - 43 | n.a. | 50.5 | 48 - 53 | n.a. |
| | $N_{D90}$ | 136 | 124 - 148 | n.a. | 161.5 | 148 - 175 | n.a. |

**Table S6:**
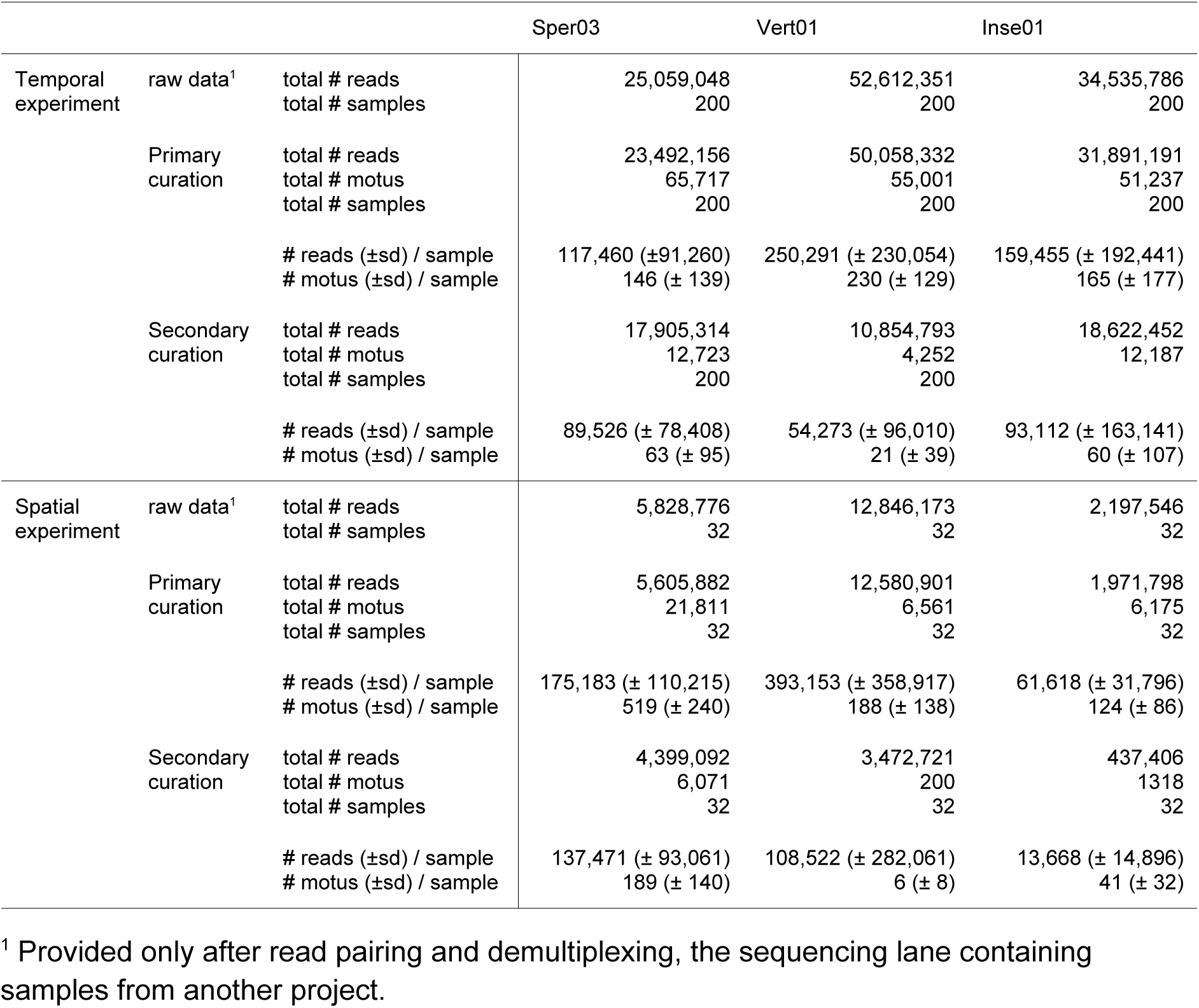
Summary statistics on raw and post-filtered data. Positive and blank controls are excluded in post-curation statistics.

## Notes

### Competing Interest Statement

The authors have declared no competing interest.

### Summary of Updates

The format of the paper has been revised to include a mode detailed introduction.

